# Arrive and wait: inactive bacterial taxa contribute to perceived soil microbiome resilience after a multidecadal press disturbance

**DOI:** 10.1101/2023.05.25.542271

**Authors:** Samuel E. Barnett, Ashley Shade

## Abstract

Long-term (press) disturbances like the climate crisis and other anthropogenic pressures are fundamentally altering ecosystems and their functions. Many critical ecosystem functions, such as biogeochemical cycling, are facilitated by microbial communities. Understanding the functional consequences of microbiome responses to press disturbances requires ongoing observations of the active populations that contribute functions. This study leverages a 7-year time series of a 60-year-old coal seam fire (Centralia, Pennsylvania, USA) to examine the resilience of soil bacterial microbiomes to a press disturbance. Using 16S rRNA and 16S rRNA gene amplicon sequencing, we assessed the interannual dynamics of the active subset and the “whole” bacterial community. Contrary to our hypothesis, the whole communities demonstrated greater resilience than active subsets, suggesting that inactive members contributed to overall resilience. Thus, in addition to selection mechanisms of active populations, perceived microbiome resilience is also supported by mechanisms of dispersal, persistence, and revival from the local dormant pool.

## Introduction

The climate crisis and anthropogenic pressure on the environment fundamentally change ecosystems and their functions. Soil microbial communities are vital to ecosystem functioning, filling important roles in biogeochemical cycling (Madsen 2011) and plant health (Mendes *et al*. 2013; Turner *et al*. 2013), among other ecosystem services. Thus, understanding the responses and functional consequences of microbial communities to severe disturbances is essential for advancing a holistic understanding of climate change and anthropogenic pressures’ impact on the natural world.

Community stability can be defined as the interplay between resistance (*i.e.,* the degree to which the community remains unchanged during a disturbance event) and resilience (*i.e., engineering resilience* (Holling 1996) or the rate at which the community returns to a pre-disturbance state) (Pimm 1984; Shade *et al*. 2012; Griffiths & Philippot 2013; Philippot *et al*. 2021). Microbial community resistance and resilience can vary substantially across soils and disturbances (Griffiths & Philippot 2013). The duration of a disturbance event, whether it is a pulse (*i.e.,* a discrete, short-term event) or press (*i.e.,* a long-term, continuous event), and whether the disturbance is compounded, may differentially affect the stability of the affected microbial community (Shade *et al*. 2012; Philippot *et al*. 2021). In light of the long-term environmental impacts of human activities and climate change across broad ecological scales, ongoing observations of microbial community dynamics to press disturbances, particularly during recovery, may illuminate the mechanisms behind community resilience to important environmental pressures.

Microbial communities tend to have low resistance to disturbances (Shade *et al*. 2012). However, the rate and degree of resilience of these communities can vary widely. The perceived variability in resilience may be due to complex, interacting, and poorly understood ecological and evolutionary mechanisms (Shade *et al*. 2012; Griffiths & Philippot 2013; Shade 2023), including the interplay of speciation, selection, and dispersal assembly processes (Vellend 2010; Nemergut *et al*. 2013). Quantifications of microbiome resilience may also be complicated by the substantial pool of dormant microorganisms that persists in many environmental systems, but especially in soils (Aanderud *et al*. 2015). Dormancy is a reversible state of extremely low metabolism that can allow bacteria to survive under poor conditions or during disturbance (Jones & Lennon 2010; Lennon & Jones 2011; Kearns & Shade 2018). Some dormant bacterial cells can remain viable for extended periods, and some can be selectively revived given conducive growth conditions (Jones & Lennon 2010; Lennon & Jones 2011). Because ecosystem response to disturbance reflects the functions of active microbiome members (Allison & Martiny 2008; Shade *et al*. 2012), it is important to account for the dynamics of active members to develop more direct predictors of ecosystem resilience and functional recovery.

The six-decade-old coal mine fire in Centralia, Pennsylvania, USA, provides a unique case study to examine the resilience of soil microbial communities to an intense, long-term heat disturbance (Shade 2018). The surface soils above the active fires are continuously hotter than ambient soil temperatures and coal combustion products percolate through the soil and open vents in the ground, altering soil chemistry at the surface (Tobin-Janzen *et al*. 2005; Janzen & Tobin-Janzen 2008; Elick 2011). The gradual advancement of the fire along the coal seams allows for direct comparisons across a gradient of disturbance intensities (*i.e.,* soil temperature) and across fire-affected, decades-old recovered, and unaffected reference soils, all within an area encompassing less than 0.35 km^2^ (Shade 2018). Furthermore, as disturbance intensity changes over time within a site, community changes can be tracked and understood in the context of the dynamic warming and cooling that define this press disturbance.

The overarching goal of this study was to understand the ecological mechanisms that support soil microbiome resilience. Our previous work in Centralia revealed that soil bacterial communities had a high capacity to recover from warming (within approximately 8-10 years) despite their multi-year exposure to the disturbance (Lee *et al*. 2017; Sorensen & Shade 2020). Advancing, we conducted a 7-year study that delineated the “whole” bacterial communities from the subset of their active members to understand the contributions of dormancy as a potential mechanism of resilience. Samples were annually collected from established Centralia sites representing fire-affected, recovered, and reference soils (Fig. 1A, Table S1). Whole bacterial communities were assessed using 16S rRNA gene amplicons, which include contributions of active, dormant, and recently deceased cells. The active subset of the whole community was determined by filtering the dataset to include only the taxa that met a conservative, population-level activity criterion based on their 16S rRNA to 16S rRNA gene ratios (Blagodatskaya & Kuzyakov 2013; Bowsher *et al*. 2019). We applied several specific methods to quantify the microbiome recovery of the whole community and their active subsets relative to the disturbance intensity, particularly time-lag and trajectory analysis (Beisner *et al*. 2003; Lamothe *et al*. 2019). Building from previous findings (Lee *et al*. 2017; Kearns & Shade 2018), we hypothesized that the microbiome resilience is largely attributable to directional changes in the active populations, indicative of environmental selection driving community turnover as the soils cool.

**Figure 1:**
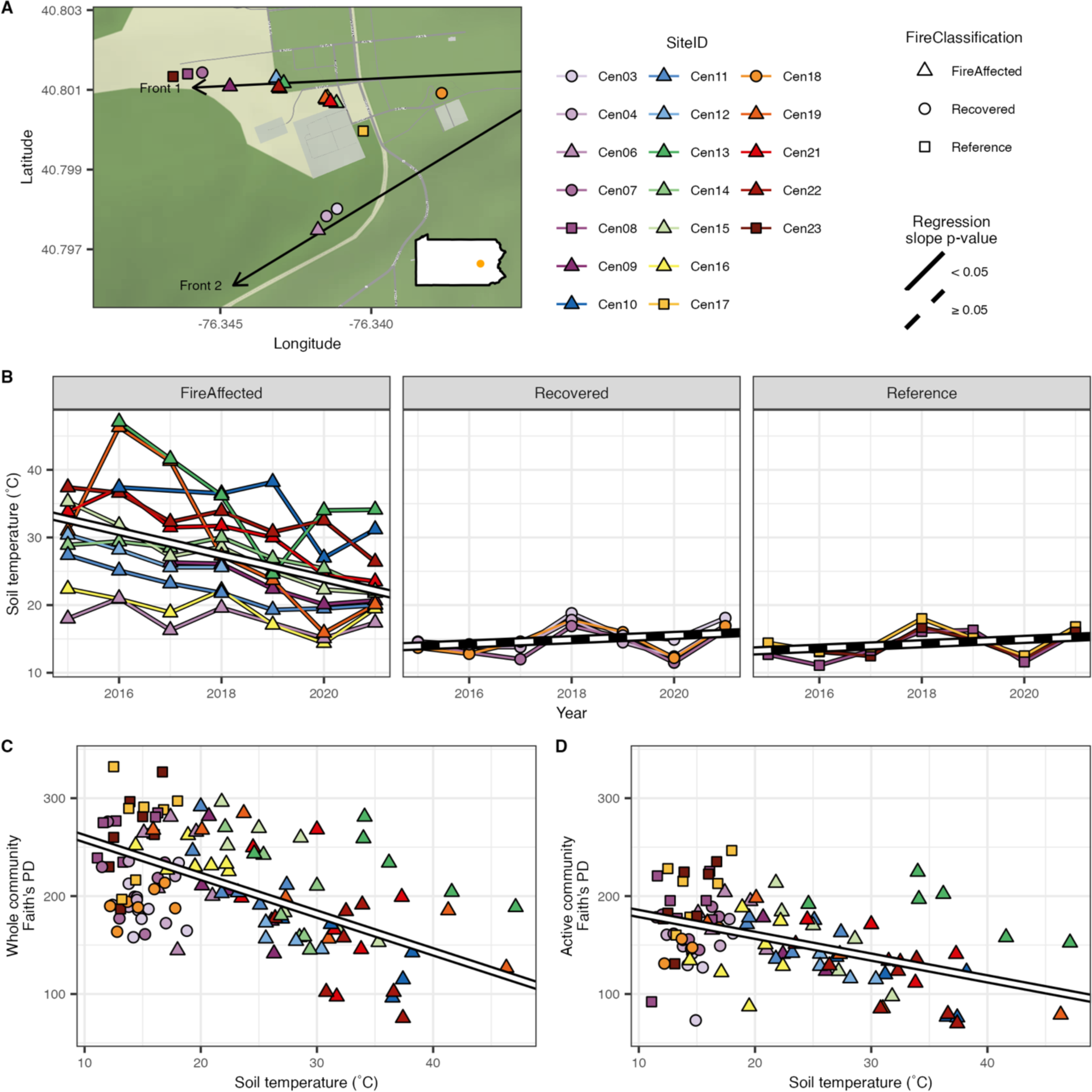
In the fire-affected sites, soils cooled over time, and along with this cooling, bacterial alpha diversity increased. A) Map of collection sites in Centralia, Pennsylvania, USA. B) Tracking of soil temperature (°C) over the seven sampling years for each site showed general soil cooling in fire-affected sites. C) The bacterial community’s phylogenetic diversity (Faith’s PD) decreased as soil core temperature increased. D) Active bacterial community subset phylogenetic diversity decreased as soil core temperature increased. For all plots, colored points and lines indicate site and fire classification. For plots B-D, white and black lines indicate linear mixed-effects regressions using site ID as the random effect, with solid white lines indicating statistically significant slopes (p-value < 0.05) and dashed lines indicating non-statistical significance (p-value ≥ 0.05). Full statistics are found in tables S2-S4, and phylogenetic diversity trends over time are found in Fig. S4.

This work illuminates the resilience of the often overlooked, inactive part of the community that is nonetheless viable and can become interactive. By considering the dormant microbial pool, this work indicates the need for careful consideration of how recovery of microbiome structure (which taxa are detected and in what relative contributions) influences microbial community functional recovery after disturbance. Thus, this study advances understanding of the complex and interacting ecological mechanisms that drive microbial community responses to disturbance.

## Materials and Methods

### Study site, soil collection, and processing

Centralia, Pennsylvania, overlies an anthracite seam that has been burning constantly since 1962 (Elick 2011). Heat and combustion gasses from the fire about 46m deep make their way to the surface, increasing surface soil temperatures and producing steam vents (Elick 2011). Nineteen sites were selected based on previous experiments (Lee *et al*. 2017; Kearns & Shade 2018), including three reference, four recovered, and 12 fire-affected (supplemental materials; Table S1). Soils were collected from each site from 2015 to 2021 in the beginning of October. 20 cm deep soil cores were collected as previously described (Lee *et al*. 2017; Kearns & Shade 2018) with sterilized PVC soil corer or trowel, sieved to 4 mm, homogenized, and aliquots for nucleotide extraction were flash frozen in liquid N_2_ (detailed in supplemental materials). Soil and air temperatures were collected with an electronic thermometer, and CO_2_ venting was measured using an EGM-5 portable CO_2_ gas analyzer (PP Systems, Amesbury, USA). Daily soil temperatures were monitored throughout the year at five sites (Table S1) using HOBO 64K pendant Temperature data loggers (UA-001-64; ONSET, Bourne USA). Soil moisture was measured gravimetrically, and soil chemistry analysis was performed by the Soil and Plant Nutrition Lab at Michigan State University (East Lansing, USA; https://www.canr.msu.edu/spnl).

### DNA/RNA extraction and processing

DNA and RNA were coextracted from 0.25 g of soil following a published phenol-chloroform protocol (Griffiths *et al*. 2000), with the alternate use of 0.1 mm zirconium bead containing BeadBug homogenizer tubes (#Z763764; Benchmark Scientific, Sayreville USA) for bead beating. Both positive (*i.e.,* mock bacterial and fungal community (Colovas *et al*. 2022)) and negative (*i.e.,* nuclease-free water) nucleotide extraction controls were included for each batch of nucleic acid extraction, and all controls were processed along with samples. DNA and RNA aliquots were processed as previously described (Lee *et al*. 2017; Sorensen & Shade 2020) and detailed in supplemental materials. To generate cDNA from RNA aliquots, DNA was degraded using the TURBO DNA-free kit (#AM1907; Invitrogen, Waltham USA) following the rigorous DNase treatment protocol for the RNA aliquot and RNA was reverse transcribed using the SuperScript III Reverse Transcriptase kit (#18080093; Invitrogen) following the standard protocol. Illumina MiSeq V4 region 16S rRNA gene amplification and sequencing of DNA and cDNA were performed by the Genomics Core Research Technology Support Facility at Michigan State University (East Lansing, MI, USA) following standard protocols as detailed in supplemental materials. Triplicate sequencing runs were performed for both DNA libraries, while single runs were performed for both RNA libraries. As detailed in supplemental materials, 16S rRNA gene counts in undiluted DNA extracts were quantified with qPCR to proxy overall Bacterial and Archaeal community sizes across the disturbance intensity gradient.

### Sequence processing

Sequence processing was performed using standard pipelines for amplicon sequencing as previously described with custom scripts (Barnett *et al*. 2020) and detailed in supplemental materials. We defined bacterial OTUs based on 16S rRNA gene sequence clusters at 97% sequence identity, including DNA and RNA sequence sets. OTUs were used rather than ASVs to avoid the potential artificial splitting of bacterial populations into multiple clusters (Glassman & Martiny 2018; Schloss 2021) and to account for potential errors that could propagate among the RNA amplicons due to the additional amplification required for reverse transcription.

### Statistical analyses

All statistical analyses were conducted using R (version 4.1.1) (R Core Team 2018). Due to the repeated measures study design, we utilized linear mixed effects (LME) models from package nlme (Pinheiro *et al*. 2020) to compare community, soil, and experimental features. In all LME models, the site was the random effect. The Benjamini-Hochberg procedure was used for all *post hoc* tests to adjust p-values for multiple comparisons. Change in soil temperature and pH over time was examined within fire classifications with LME models. Differences in soil pH across fire classifications were tested using ANOVA and post hoc Tukey’s.

OTU counts from triplicate libraries were summed for DNA samples. Samples with less than 100,000 reads were removed because these were clear outliers regarding sequence output, suggesting low quality. Sites with less than three years of adequate DNA sequencing were removed, and the resulting OTU table was rarefied to 161,171 reads and used for the whole community analysis. To construct an OTU table for the active community, DNA samples without a corresponding RNA sample after quality filtering and sites with less than three years of paired DNA and RNA samples were removed, and the resulting OTU table was rarefied to 100,491 reads. OTUs found in an RNA sample but not the corresponding DNA sample (*i.e.,* phantom OTUs, Fig. S1, supplemental material) were given a count of 1 in the DNA sample (Blagodatskaya & Kuzyakov 2013; Bowsher *et al*. 2019). OTUs with an RNA: DNA ratio ≥ 1 in each paired RNA and DNA sample were then defined as active (Blagodatskaya & Kuzyakov 2013; Bowsher *et al*. 2019). Analysis of the active community subset used the OTU tables from the filtered DNA samples to only include active OTUs (*i.e.,* zero count wherever DNA and cDNA count was zero or RNA: DNA ratio was < 1). Critically, DNA relative abundances (not cDNA relative abundances) were used to determine active taxon dynamics, as the relativized abundances of 16S rRNA are not indicative of population abundances due to the known individual- and population-level heterogeneity in gene regulation, ribosomal gene copy number, post-translational modification, ribosome stability, etc. (Blazewicz *et al*. 2013).

Alpha diversity was measured as Faith’s phylogenetic diversity index (PD). Change in PD over time within fire classifications and across soil temperature regardless of fire classification was examined with LME models. Beta diversity was measured using abundance-weighted UniFrac distance calculated with package phyloseq (McMurdie & Holmes 2013). Variation in community structure across fire classification, sampling year, and their interaction was performed using PERMANOVA blocked by site ID. *Post hoc* analysis of variation in community structure across fire classifications within each year or across years within each fire class was also performed using PERMANOVA blocked by site ID. Variation in community structure explained by edaphic factors and temperature was examined using a constrained correspondence analysis with package phyloseq. Selection of edaphic factors included in the final models was accomplished with function ordistep from the vegan package (Oksanen *et al*. 2018) with the initial full model incorporating soil temperature, CO_2_ emission, pH, percent soil organic matter, and phosphate, potassium, calcium, magnesium, iron, nitrate, ammonium, sulfate, and arsenic concentrations.

We used two methods to assess community change over time. First, time-lag analysis measures the distance between communities within a site over increasing time-steps. The shape, particularly the slope, of the relationship between community distance and time-lag can indicate directional shifts in communities during recovery (Lamothe *et al*. 2019). Time-lag analysis was performed separately for each fire classification using LME models. Abundance-weighted UniFrac distance across time points within a site was the response variable, and the square root of time point difference was the fixed effect. Slopes of time-lag linear regressions for each site were then regressed to the maximum soil temperature at each site, regardless of fire classification, using a linear model. Second, community trajectory analyses measure directionality in community structure over time, which can indicate community recovery (Lamothe *et al*. 2019). Trajectory directionality was calculated using package ecotraj (De Cáceres *et al*. 2019; Sturbois *et al*. 2021) and was compared across fire classifications using a Kruskall-Wallis test and *post hoc* pairwise Dunn’s tests. Trajectory directionality was also regressed to maximum soil temperature at each site, regardless of fire classification, using a linear model. To further assess whether the bacterial communities were becoming more like the undisturbed reference communities as soils cooled, we tested whether the distance in community structure between disturbed soils (*i.e.,* fire-affected and recovered soils) and reference soils within years was associated with time or disturbed soil temperature. We used LME models to compare the mean abundance-weighted UniFrac distance between the disturbed site and all three references, paired by timepoint, to year or soil core temperature.

We calculated the percentage of the Bray-Curtis dissimilarity within each site attributed to active taxa using previously described methods (Shade *et al*. 2014), see Supplemental Methods. To examine the influence of OTUs that become inactive or newly active over time, we separately analyzed OTUs active in the first timepoint of each timepoint pair (*e.g.,* 2016 when comparing 2016 to 2020) and in the second timepoint of each pair (*e.g.,* 2020 when comparing 2016 to 2020). The percentage of the Bray-Curtis dissimilarity attributed to active taxa was compared across fire classifications using Kruskall-Wallis and *post hoc* pairwise Dunn’s tests and across time differences between samplings within each fire class using LME models.

Upset plots were made using custom code based on the R package UpSetR (Conway *et al*. 2017), see Supplemental Methods. Community assembly processes contributing to the whole bacterial communities’ structure were accomplished using NTI calculations (Stegen *et al*. 2012, 2015; Dini-Andreote *et al*. 2015; Barnett *et al*. 2020; Ning *et al*. 2020), see Supplemental Methods. βNTI values for site pairs within each fire classification were compared across these classes using Kruskall-Wallis and *post hoc* pairwise Dunn tests. The relationship between βNTI values for site pairs and differences in soil temperature, the difference in soil pH, and their interaction were examined using a linear model after scaling and centering both factors to make them directly comparable.

## Results

### Soil temperature and pH vary over fire classification

As expected, fire-affected soils were generally hotter than recovered and reference soils. Notably, the fire-affected soil temperatures decreased over the seven years, yet no consistent changes over time were observed in recovered or reference sites (Fig. 1B, Table S2; LME models). This pattern was confirmed by daily temperatures in selected sites that were hourly tracked over the seven years by the *in situ* HOBO monitors (Fig. S2). Soil pH did not change over time (Fig. S3A, Table S2; LME models), though it did overall differ across fire classifications (ANOVA: DF = 2, F = 14.54, p-value < 0.001). pH was lower in recovered sites than either reference or fire-affected sites, and the pH range across fire-affected sites was large (Fig. S3B, *post hoc* Tukey’s: p-values < 0.05).

### Bacterial diversity changes across fire classification and over time

Faith’s phylogenetic diversity (PD) increased over time in fire-affected soils for both whole communities and active subsets but no increase in recovered or reference soils (Fig. S4; Table S3; LME models). When time and fire classification were ignored, PD increased as soil temperature decreased for whole communities and active subsets (Fig. 1C and D, Table S4; LME models). Overall bacterial and archaeal counts, as determined by qPCR, showed no significant relationship to soil temperature across our soils (Table S5; LME model).

For whole bacterial communities, phylogenetic beta diversity (weighted UniFrac) was explained by fire classification (18.0% variation explained), time (5.7%), and their interaction (4.0%) (Fig. S5, Table S6; PERMANOVA). *Post hoc* tests within each fire class showed that time was explanatory only in fire-affected sites (Fig S6, Table S7). While fire classification was explanatory in all years, the variation explained (R^2^) decreased between 2015 and 2018 (Fig. S7, Table S7). Differences across fire classifications could be attributed to differences in beta-dispersion (ANOVA; df = 2, F = 34.722, p-value < 0.001). Specifically, dispersion was higher among fire-affected soils than recovered and reference soils (Table S8).

For the active subset of the community, fire classification (12.4% variation explained) and time (5.7%), but not their interaction, explained phylogenetic beta diversity (Fig. S8, Table S6; PERMANOVA). Beta-dispersion again varied across fire classifications for the active subset (ANOVA; df = 2, F = 17.536, p-value < 0.001). Dispersion across fire-affected soil was higher than recovered and reference soils, though the effect was smaller than for the whole community (Table S8).

Measured edaphic factors, particularly soil temperature, pH, and pH-associated properties (*e.g.,* Ca, Mg, and Fe concentration), explained a large portion of the variation in community structure for both whole communities (Fig. S9A; variation explained = 45.342%) and active subsets (Fig. S9B; variation explained = 35.789%). As expected, the variation between fire-affected and both reference and recovered sites was largely associated with soil temperature. In contrast, variation across and within the reference and recovered sites was primarily associated with pH (Fig. S9).

### Disturbance intensity drives community trajectories

For both the whole bacterial communities and active subsets, time-lag analysis showed increasing community dissimilarity as the timespan between the samplings increases for fire-affected sites but no relationship for either recovered or reference soils (LME models; Fig. 2A and B, Table S9). The slopes of the time-lag relationships were positively correlated to the maximum soil temperature of the sites (Fig 2C and D; linear regression; whole community: R^2^ = 0.67, p-value < 0.001, active subset: R^2^ = 0.30, p-value = 0.011). Notably, the relationship between the time-lag slope and maximum soil temperature was stronger for the whole community than the active subset.

**Figure 2:**
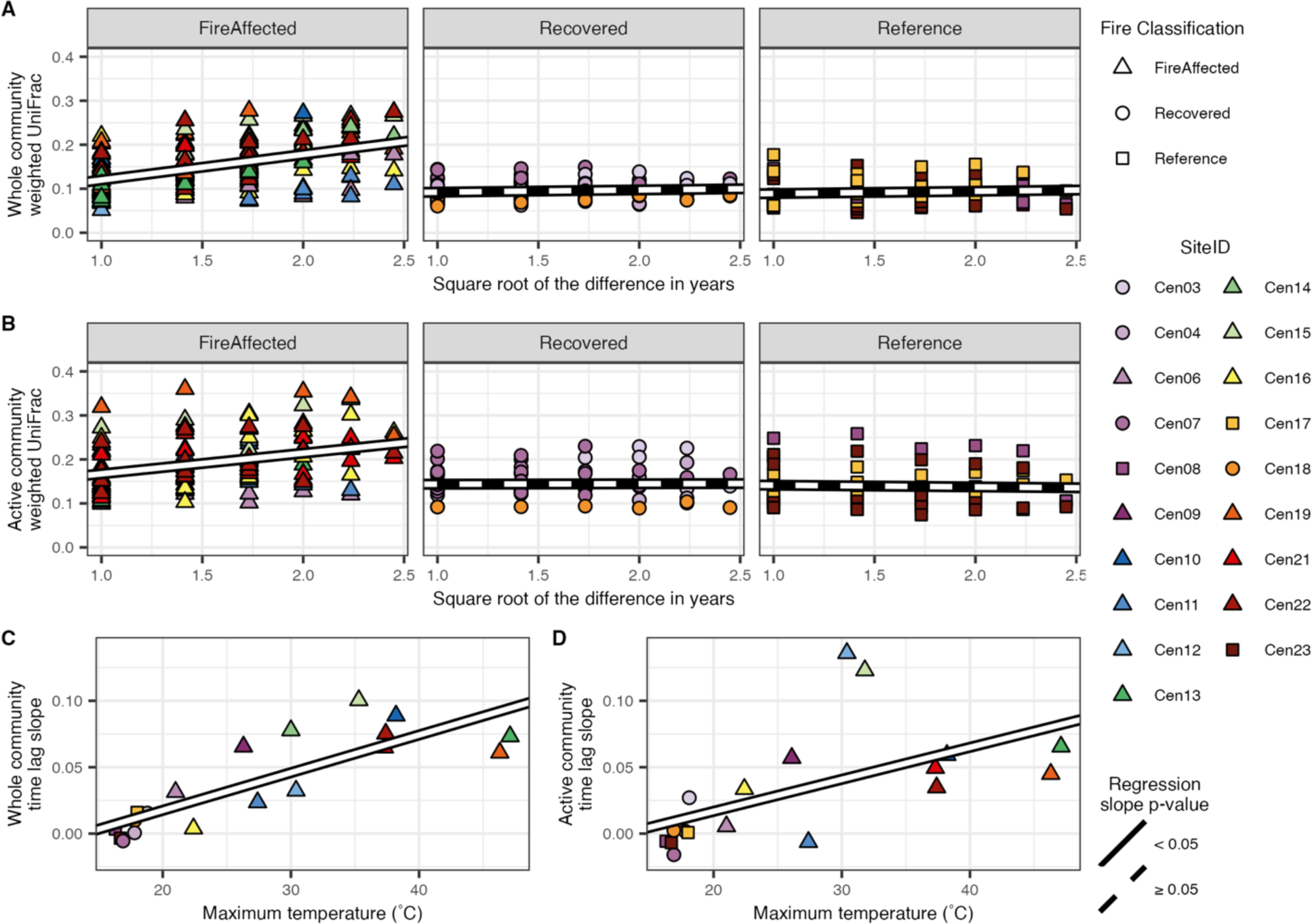
Time-lag analysis demonstrated that variation in the whole bacterial community and active subset structure between samplings of the same site increased as the time difference between sampling increased. However, this trend was only statically significant for fire-affected sites. A) Whole community time-lag analysis. B) Active community subset time-lag analysis. C) Slopes of whole community time-lag regressions made individually for sites increased as maximum soil temperature (proxy of disturbance intensity) increased across sites. D) Slopes of active community subset time-lag regressions made individually for sites increased as maximum soil temperature increased across sites. Points indicate site ID and fire classification. White and black lines indicate linear mixed-effects regressions using site ID as the random effect (A and B) and linear regression (C and D) with solid white lines indicating statistically significant slopes (p-value < 0.05) and dashed lines indicating non-statistical significance (p-value ≥ 0.05). Full statistics for the linear mixed effects regressions are provided in Table S8.

For both the whole communities and active subsets, there was higher directionality in the fire-affected sites than either recovered or reference sites, with no significant difference between the latter classifications (Fig. 3A and B; Kruskal-Wallis rank sum test; whole community: chi-squared = 9.04, df = 2, p-value = 0.01, active subset: chi-squared = 8.67, df = 2, p-value = 0.01; *post hoc* Dunn’s tests). Community directionality was positively correlated with maximum soil temperature (a proxy for disturbance intensity), with, again, a stronger relationship for the whole communities relative to the active subsets (Figs. 3C and D; linear regression; whole community: R^2^ = 0.55, p-value < 0.001, active subset: R^2^ = 0.20, p-value = 0.036).

**Figure 3:**
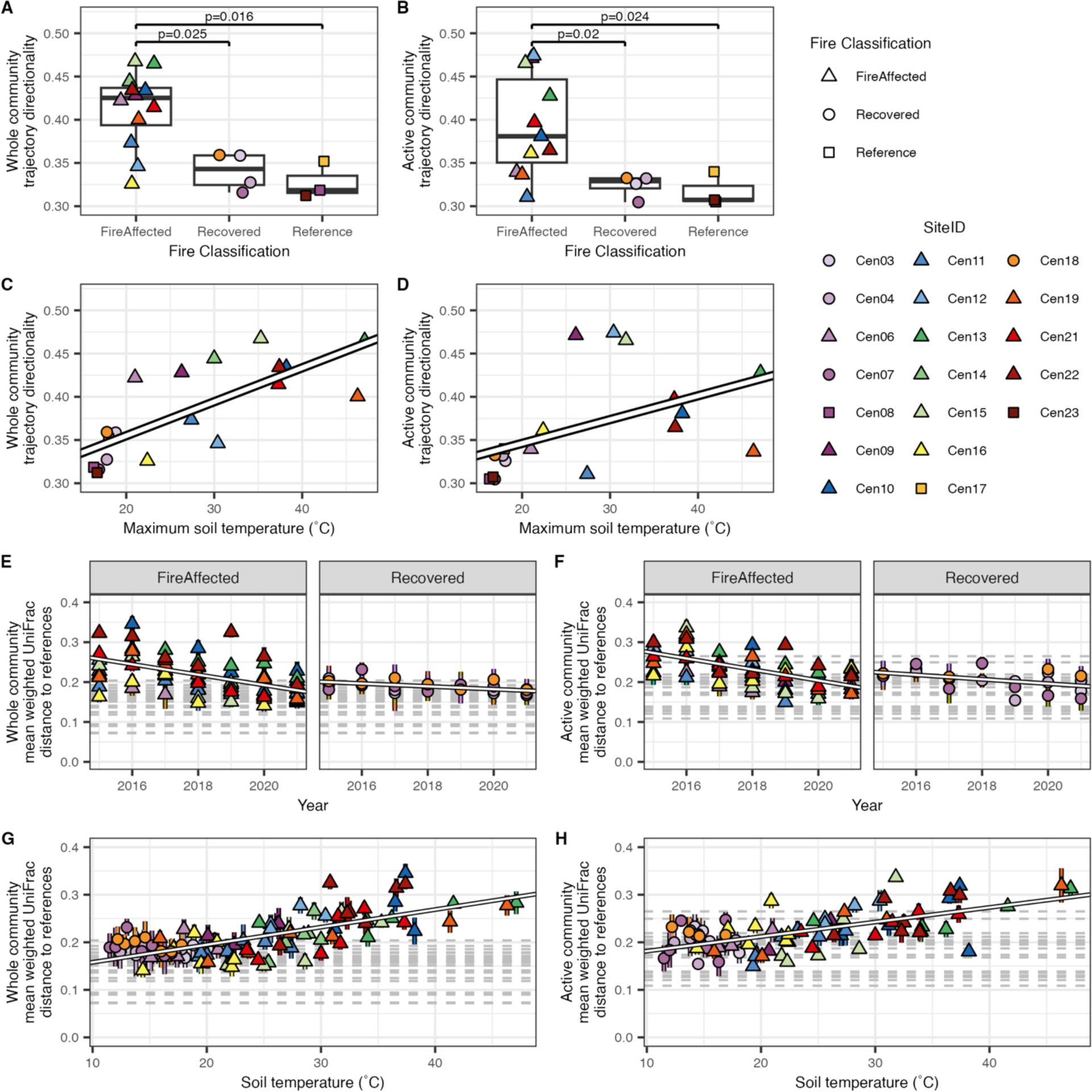
Trajectory analysis demonstrated that directionality in both whole bacterial community (A) and active subset (B) structural shifts over time were generally highest in fire-affected soils compared to recovered and reference soils. Post-hoc pairwise Dunn’s test p-values < 0.05 are displayed above statistical significance indicator bars. Directionality in both whole bacterial community (C) and active subset (D) structural shifts increased as maximum soil temperature within sites increased. For both whole bacterial communities and active subsets, disturbed soil communities became more structurally similar to the undisturbed reference communities over time as soils cooled (E and F, respectively) and soils cooled (G and H respectively). Structural dissimilarity was calculated as the mean weighted UniFrac distance between fire-affected and recovered sites and the three reference sites paired within the sampling year. White lines indicate linear regressions (C and D) and linear mixed-effects regressions using site ID as the random effect (E-H), which had statistically significant slopes (p-value < 0.05). For figures A-D, points indicate all samples’ site ID and fire classification. For figures E-H, points indicate the site ID and fire classification (fire affected or recovered) of the disturbed site with error bars representing the standard error for the distance between the sample and the three paired reference samples from the same year. For E-H, grey dashed horizontal lines indicate community distances between the reference sites within each year.

We next tested whether the change in community structure between disturbed soils (*i.e.,* fire-affected and recovered soils) and reference soils within years changed over time or was associated with soil temperature. We found that for both whole communities and active subsets, disturbed communities tended to become more like the reference communities over time and as temperature decreased (Fig. 3E-H; LME models). While fire-affected and recovered soils demonstrated increased similarity to reference communities over time, the slope of the change in community distances was higher for fire-affected communities than recovered communities (Fig 3E-F).

### Active members contribute little to shifts in the whole community

Across most sites, less than 50% of the Bray-Curtis dissimilarity in the whole community was attributable to active OTUs. For fire-affected sites, the dissimilarity attributed to active taxa decreased as the period between sampling years increased (Fig. 4, Table S10; LME models; p-values < 0.05). This trend was statistically supported when partitioning taxa as active at the earlier and later time points in the year-pairing. In contrast, no such trend was observed for recovered or reference sites.

**Figure 4:**
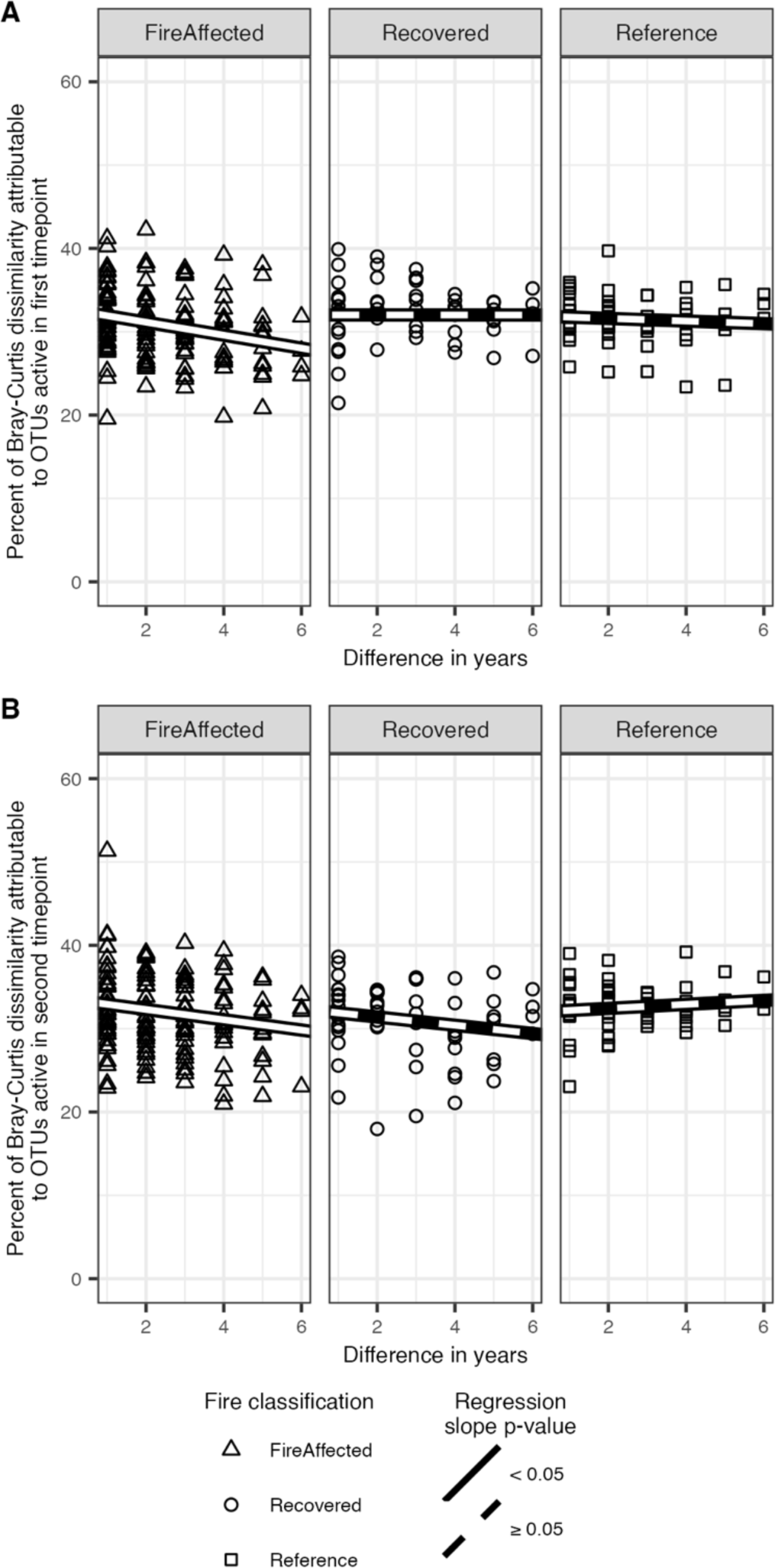
For fire-affected sites, the whole bacterial community structural dissimilarity attributable to the active subset decreases as the time between sampling time points increases. In other words, less of the whole community dissimilarity is attributable to the active subset across larger timescales for fire-affected sites but not for recovered or reference sites. This holds across site pairs for OTUs active in (A) the early timepoint (*e.g.,* 2016 when comparing 2016 and 2020) and in (B) the later timepoint (*e.g.,* 2020 when comparing 2016 and 2020). Bray-Curtis dissimilarity within a site was used to partition OTUs active in either each pair’s early or late time points. White and black lines indicate linear mixed-effects regressions using site ID as the random effect, with solid white lines indicating statistically significant slopes (p-value < 0.05) and dashed lines indicating non-statistical significance (p-value ≥ 0.05).

### Interannual OTU activity dynamics vary across fire-affected and unaffected soils

Many OTUs had distinctive yearly activity patterns (Fig. 5 and Figs. S10 and S11). For fire-affected sites (Fig. 5A), most OTUs tended to have consistent activity in later sampling years (*e.g.,* 2018-2021) but not earlier (*e.g.,* 2015-2017). This trend suggests that many OTUs became active in later years for fire-affected sites, corresponding to cooling soil temperatures and recovery. For reference and recovered soils (Fig. 5B), most OTUs had no clear annual progression or delineation, suggesting more random activity patterns in these soils that experienced ambient temperatures. Activity groups were taxonomically diverse for both cases, with no obvious enrichment in taxa at the phylum level or *Proteobacteria* class.

**Figure 5:**
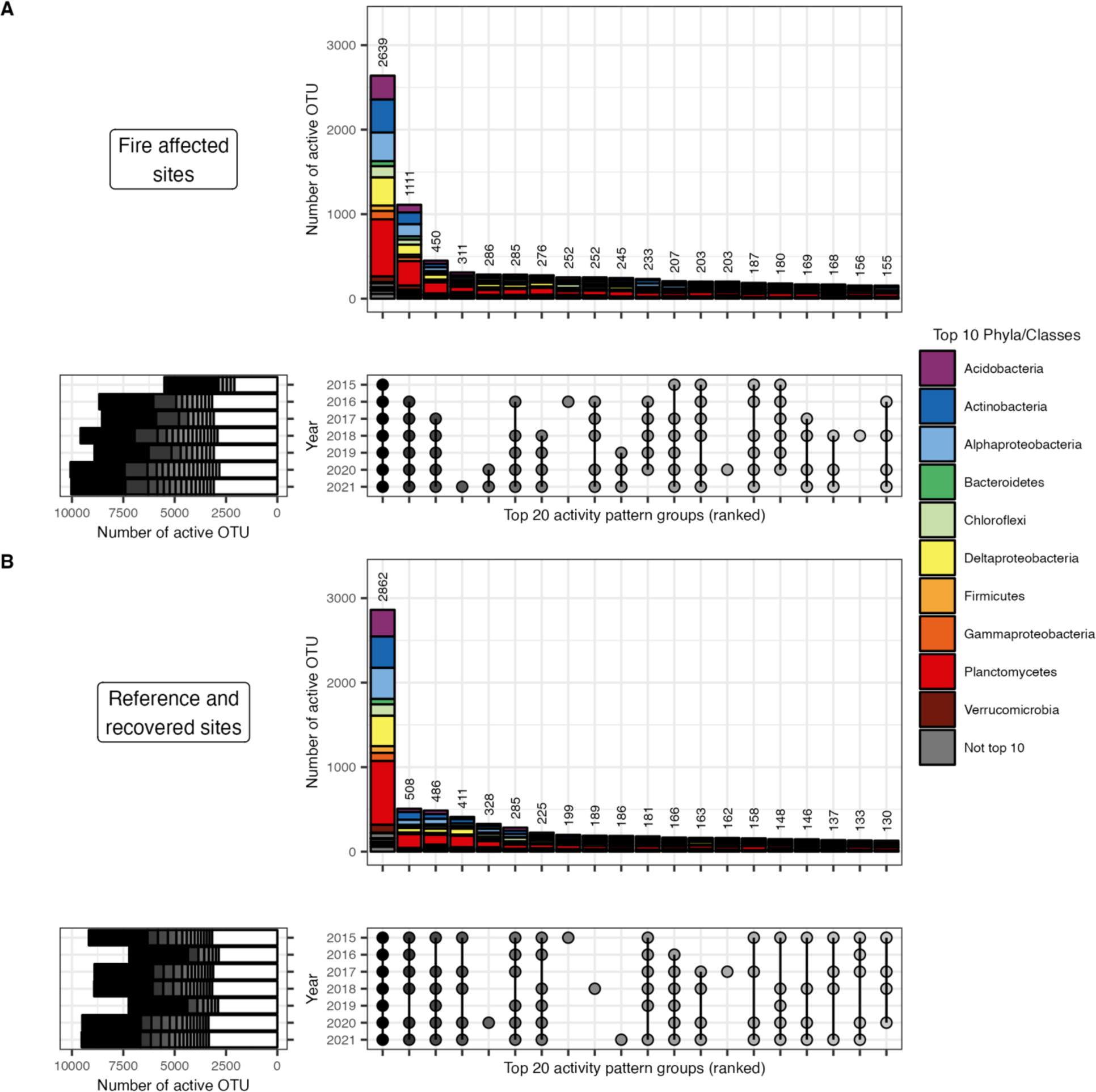
UpSet plots of the top 20 (as determined by OTU count) observed activity dynamics across years for (A) fire-affected sites and (B) combined recovered and reference sites. All sites sampled each year were combined such that OTUs were defined as active if they had an RNA: DNA ≥ 1 in at least one site within that year. The bottom right plots show the activity patterns of the top 20 groups, with dots indicating activity within that year and groups ordered by OTU membership. The top vertical bar chart shows the number of OTUs within each activity group, colored by the top 10 phyla (class for *Proteobacteria*). The bottom left horizontal bar chart shows the number of active OTUs within each year, colored by their membership in the top 20 activity groups. White sections of the bars indicate OTUs from the other 107 groups (see Figs. S10 and S11 for all 127 activity groups). For the fire-affected sites (A), most of the top 20 activity groups do not have detectable activity in the first few sampled years, while most of the top 20 activity groups for reference and recovered sites (B) have detectable activity during those years.

### Community assembly processes for whole communities differ across fire classifications

While the majority of within-year, site-to-site comparisons indicate dominance of homogeneous selection (βNTI < -2), phylogenetic turnover among fire-affected sites tended to be higher than among recovered or reference sites (Fig. 6A; Kruskal-Wallis rank sum test: DF = 2, chi-squared = 68.399, p-value < 0.001; post hoc Dunn test) with many site pairs demonstrating weak selection (-2 ≤ βNTI ≤ 2) or variable selection (βNTI > 2). This pattern was consistent across seven years (Fig. 6D and S12). For these site pairs with weak selection, homogenizing dispersal (RC_bray_ < -0.95), followed by drift alone (-0.95 ≤ RC_bray_ ≤ 0.95), tended to be the dominant assembly process (Fig. 6D). Scaled between-site differences in soil pH and temperature were both positively associated with βNTI, but their interaction was not (Fig. 6B and C, Table S11; linear regression: R^2^ = 0.5031, p-value < 0.001). The coefficient for scaled difference in pH (2.63) is much larger than that of scaled difference in temperature (0.42).

**Figure 6:**
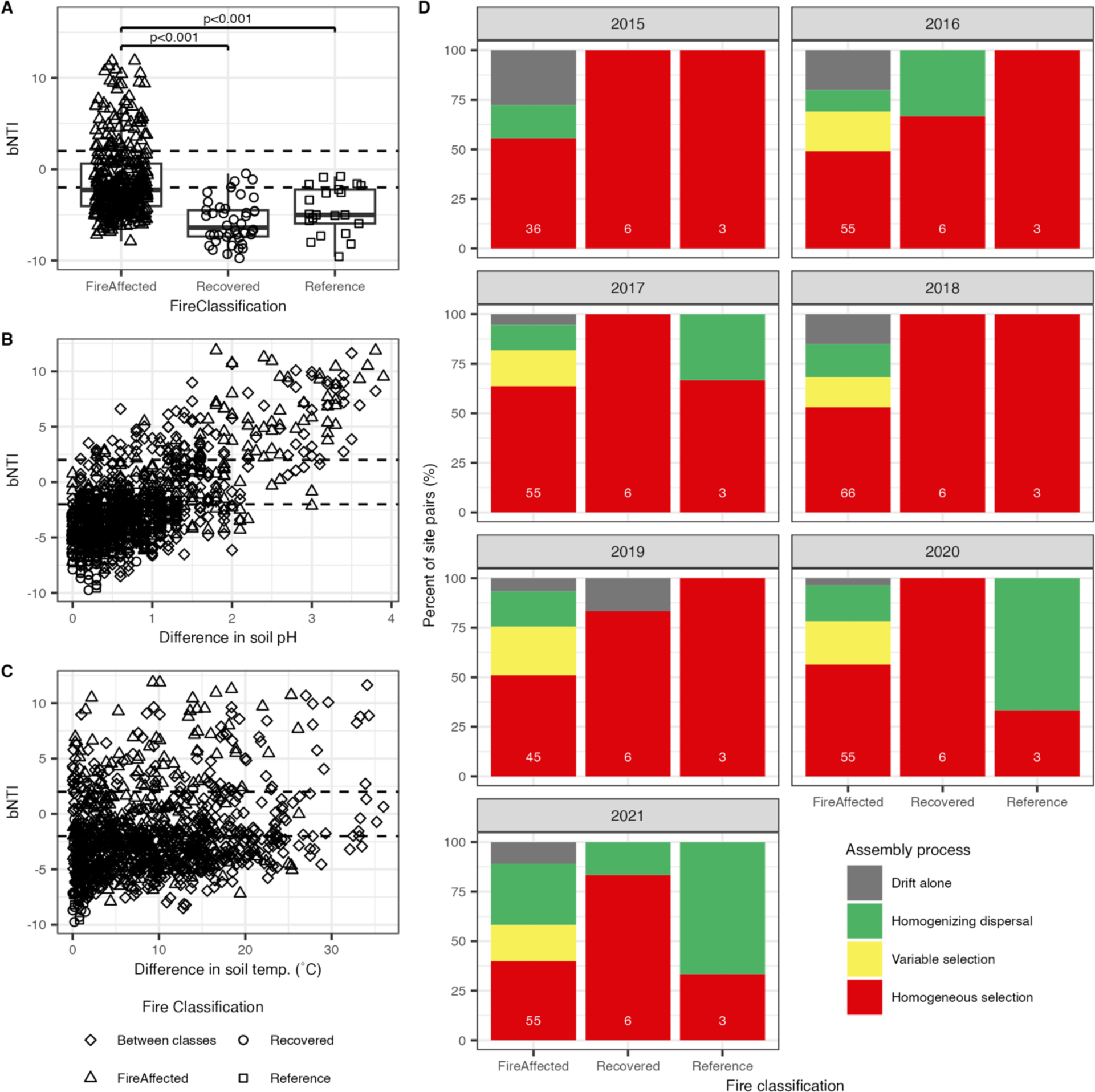
The whole bacterial community assembly dynamics across sites were measured with standardized phylogenetic and structural turnover. A) βNTI is generally higher between fire-affected sites than recovered or reference sites whose site pairs almost always have a βNTI < 2, indicating homogeneous selection. Post hoc pairwise Dunn’s test p-values < 0.05 are displayed above statistical significance indicator bars. B) Regardless of fire classification, βNTI tended to increase as the difference in soil pH between sites increased, indicating pH-related selection pressures in these soils. C) Regardless of fire classification, βNTI tended to increase as the difference in soil temperature between sites increased, indicating some temperature-related selection pressures in these soils. Linear regressions, including these two relationships and their interaction, are provided in Table S10. D) Percentage of site pairs within each fire classification and year for which community assembly can be attributed to the five assembly processes examined with βNTI and RC_bray_. The numbers of site pairs are indicated by white numbers at the bottom of each bar. All site pairs were within each year, not across years.

## Discussion

We used long-term soil microbiome surveys from Centralia, Pennsylvania, USA, to examine the resilience of whole bacterial communities and their active members impacted by heat press disturbance from a subterranean coal seam fire. Over seven years, we observed general cooling of the fire-affected sites, indicating a gradual reduction in disturbance intensity. Cooling was not unexpected at these locations (Elick 2011). While heat intensity decreased in most sites, other fire-induced environmental impacts continued, exemplified by elevated local CO_2_ concentrations and steam venting observed in some soils only a few degrees Celsius above ambient temperature. The gradual change in disturbance intensity provided a unique opportunity to observe the transition and recovery of the microbial communities during both early stages and later stages of recovery (*i.e.,* cooling fire-affected and recovered sites, respectively). Temporal trends in bacterial diversity revealed strong evidence for the resilience of the microbiomes in both the whole communities and their active subsets. In comparison, communities in recovered and reference sites were more stable over the years. These findings agree with our previous study, which included soils compared along the fire-impact gradient within one sampling year, that the whole bacterial communities in Centralia soils are resilient (Lee *et al*. 2017).

Dispersal is an important microbiome assembly mechanism for soil (Albright & Martiny 2018; Barnett *et al*. 2020; Evans *et al*. 2020; Walters *et al*. 2022). Across Centralia sites, when selection was weak, homogenizing dispersal was the dominant assembly process. Dispersal likely explains the increased alpha diversity observed as fire-affected soils cool. However, even when dispersal is high, immigrant taxa may not necessarily be immediately active or later become established. Rather, their DNA may remain extractable either in intact cells (dormant or dead) or as extracellular DNA (Morrissey *et al*. 2015; Carini *et al*. 2017; Lennon *et al*. 2018). The microbial conveyor belt hypothesis proposes that dormancy aids microbial dispersal capability (Mestre & Höfer 2021), though more supporting evidence is needed (Choudoir & DeAngelis 2022). During a 45-week mesocosm experiment with soil from Centralia reference sites that explicitly controlled dispersal (Sorensen & Shade 2020), we observed that post-disturbance activation of dispersed taxa contributed to overall resilience. Building from these collective results, we present an updated working hypothesis to integrate dispersal and dormancy mechanisms in soil microbiome resilience. We hypothesize that dispersed members from the regional species pool progressively replace taxa lost to disturbance as thermal selection weakens, but some likely maintain dormancy while conditions remain suboptimal. This process effectively rebuilds and accumulates the dormant “seed bank” but does not contribute immediately to functional recovery (*i.e.,* active populations) until conditions become favorable for activity. Thus, re-establishing the microbial seed bank during or after a disturbance could provide a restoration management strategy (Moreira-Grez *et al*. 2019; King & Bell 2022; Shade 2023).

Patterns of OTU activity over time in fire-affected sites demonstrated that many bacteria are likely to be activating (either after dispersal or from the local seed bank) in the fire-affected soils over the last few years as the soils have cooled. Of the top 20 interannual activity patterns, most taxa tended to have observed activity only in the later sampling years. In other words, many active taxa become detectably active after one to three years of no detectable activity in fire-affected soil. This is best exemplified by the second most numerous activity pattern that includes activity detected in 2016-2021 but not 2015, possibly suggesting significant activation of taxa after this first sampling year in particular. In reference and recovered sites, the top 20 patterns exhibited random years in which no activity is detected, including patterns with detectable activity in the first few sampling years. We further see that the most numerous activity patterns include many phyla, suggesting that activation and perhaps recovery of the active community subset is phylogenetically indiscriminate at high taxonomic levels.

Notably, the temporal changes within the whole communities were greater than in their active subsets, indicating differences in perceived resilience and recovery between the active and whole communities. For example, time explains more variation in beta diversity for whole communities than for active subsets, and no interaction was found between time and fire classification for active communities. Similarly, the relationships between maximum temperature and both time lag slopes and trajectory directionality were weaker for active subsets than the whole communities. When considering variation in the whole community structure within a site over time, active OTUs tended to explain a smaller percentage of the community dissimilarity as the timescale increased, but only for fire-affected sites. This important finding indicates that over multiple years, changes in the structure of the active subset account for less of the change in overall community structure in recovering soils. Thus, we propose that while the overall bacterial community is largely resilient, the active populations collectively may recover slower. This conclusion refutes our original hypothesis that bacterial community resilience is largely attributable to the dynamics of active populations. It suggests that dispersal enables bacterial taxa from undisturbed soils to arrive and persist in the fire-affected soils as they cool, with some immigrants remaining dormant though persistent as disturbance intensity slowly wanes. This accumulation of inactive microbial diversity throughout a disturbance may enable recovery of the bacterial community after the disturbance; this should be investigated in future studies.

pH and associated edaphic properties are well-known drivers of bacterial community diversity and soil assembly (Fierer & Jackson 2006; Malik *et al*. 2017; Tripathi *et al*. 2018; Barnett *et al*. 2020; Yavitt *et al*. 2020). Many pH related properties explained structural differences among Centralia communities (e.g., Ca, Mg, Fe, etc.) and βNTI (Stegen *et al*. 2012, 2015) correlated to difference in soil pH, together indicating pH-related selection across Centralia soils. βNTI also correlated to difference in soil temperature, though this relationship was weaker than that of soil pH, and there was no statistically significant interaction between these factors. We thus propose that while soil pH selection acts upon Centralia soil bacterial communities, soil heating imposes either alternate selection pressures or increased stochasticity. Soil acidification may be a legacy effect of mining (Rice & Herman 2012; Wang *et al*. 2020) or coal fires in this area (Tobin-Janzen *et al*. 2005; Janzen & Tobin-Janzen 2008; Lee *et al*. 2017) and may promote alternative stable states among the recovered communities distinct from unaffected soils (Quadros *et al*. 2016).

Other mechanisms may also have contributed to the patterns we observed. First, cyclical seasonal disturbance intensity patterns, as observed from HOBO temperature monitors, may produce temporary conditions suitable for the growth or maintenance of fire-sensitive populations (Philippot *et al*. 2021). Second, as soils are extremely diverse (Rondon *et al*. 1999), the non-detection of rare taxa is a distinct possibility despite our deep sequencing, as demonstrated by the rarefaction curves. Third, extracellular DNA (eDNA) may contribute to the divergence in the whole community and active subset dynamics between fire-affected and unheated soils because eDNA in soil degrades more rapidly at higher temperatures (Sirois & Buckley 2018). However, a large influence of eDNA on the observed interannual patterns is not expected due to the short lifespan of nucleic acids in soils in general.

Broadly, our findings and working conceptual model for bacterial community recovery from press disturbance have implications for understanding ecosystem recovery to large, long-term stress. We interpret the observed relatively high resilience of the whole community with somewhat lower resilience of the active subset to indicate that, while microbial community functional potential may recover rapidly along with the demographic recovery of members lost to the disturbance, the realized activity of these communities may take longer to recover because it is harbored within the dormant pool. As only the active subset of the community performs ecosystem functions and services, it is likely that biologically mediated soil processes, such as biogeochemical cycling, may remain altered for some period after disturbance intensity wanes, especially for functions with low redundancy among populations. If ecosystem function is used to measure ecosystem recovery (Harris 2003), then whole bacterial community surveys will provide a biased view of the recovery progress. However, our findings provide a positive outlook on the recovery potential of disturbed soils. Understanding the stability and potential for recovery of microbial communities will help to predict changes to vital ecosystem functions and develop methods of microbiome restoration.

## Statement of authorship

AS conceptualized the research, AS and SB collected soil samples at the research site (investigation), SB performed laboratory work and conducted formal analysis (investigation), SB performed data curation, SB wrote the original draft manuscript, SB and AS contributed to review and editing of the manuscript, SB performed data visualization, and AS provided supervision, administration, and funding acquisition.

## Data accessibility statement

Raw sequences are available on the NCBI short read archive BioProject PRJNA973689. Representative OTU sequences in FASTA format, along with metadata, OTU table, taxonomy table, and phylogenetic tree, all in a phyloseq object, are available through figshare DOI: 10.6084/m9.figshare.23060354. Sequence processing and analysis code are available through GitHub at https://github.com/ShadeLab/Centralia_RNA_DNA_multiyear.

## Acknowledgments

We thank Tammy Tobin-Janzen for Centralia research consultation and local sampling support and Keara L Grady for coordinating fieldwork logistics and organization of multi-year soil procurement and processing. We additionally thank SH Lee, JW Sorensen, TK Dunivin, JL Chodkowski, N Stopnisek, and M Mechan Llontop for collecting samples. This work was supported by the U.S. National Science Foundation CAREER award #1749544 to AS. AS acknowledges the French Centre National de la Recherche Scientifique (CNRS) support. We declare no conflicts of interest.

