## supplemental for "Arrive and wait: inactive bacterial taxa contribute to perceived soil microbiome resilience after a multidecadal press disturbance"

*This file contains the following sections:*

1. Supplemental Materials and Methods
2. Supplemental Figures
3. Supplemental Tables

### 1. Supplemental Materials and Methods

*Soil collection and processing*

Based on previous experiments, we selected 23 sites along the two main fire fronts (Lee *et al.* 2017; Kearns & Shade 2018). Soils were collected from each site from 2015 to 2021 in the first or second week of October. Four sites were removed from the experiment before analysis due to exhaustion of available soil, a landslide, or exceedingly low DNA concentrations in the collected soil.

Where possible, 20 cm deep and 5.1 cm in diameter soil cores were collected using ethanol-sterilized PVC soil corers. When coring was impossible, primarily in vents, the soil was collected with an ethanol-sterilized trowel. Collected soil was homogenized and sieved to 4 mm. Soil for nucleic acid extraction was immediately transferred to a sterile 50 ml falcon tube and flash-frozen in liquid $N_2$. The time from soil collection to flash freezing was generally under 4 minutes. Flash-frozen soils were stored at -20˚C or on ice while transported to the lab and then stored at -80˚C until nucleic acid extraction. The soils used for soil chemistry analyses were transferred to a Whirl-Pak bag and stored on ice or at 4˚C while transported to the lab. Soil moisture was measured gravimetrically with ~10 g of soil. The Soil and Plant Nutrition Lab at Michigan State University (East Lansing, USA; https://www.canr.msu.edu/spnl) processed all other soil chemistry analyses according to standard protocols. Measurements included pH, concentrations in ppm of phosphate, potassium, calcium, magnesium, iron, nitrogen from nitrate, nitrogen from ammonium, sulfur from sulfate, arsenic, and percentage of organic matter. Just before soil collections, soil core temperature and air temperatures were measured using ethanol-sterilized electronic thermometers, and $CO_2$ venting measurements were measured using an EGM-5 portable $CO_2$ gas analyzer (PP Systems, Amesbury USA). Daily soil temperatures were monitored at five sites (Table S1) using HOBO 64K pendant Temperature data loggers (UA-001-64; ONSET, Bourne USA) buried ~10cm below the surface near the coring site and recovered each year.

*DNA/RNA extraction and processing*

DNA and RNA were coextracted from 0.25 g of soil following the Griffiths et al. phenol-chloroform protocol (Griffiths *et al.* 2000), with the alternate use of 0.1 mm zirconium bead containing BeadBug homogenizer tubes (#Z763764; Benchmark Scientific, Sayreville USA) for bead beating. DNA and RNA aliquots of extracted nucleic acid were stored at -80˚C for further processing. DNA and RNA processing was performed within a month of extraction to prevent excessive RNA degradation. DNA concentration was determined using Qubit dsDNA BR Assay Kit (#Q32853; Invitrogen, Waltham USA). If possible, DNA aliquots were diluted to ~2 ng/µl DNA, and RNA aliquots were diluted to ~200 ng/µl DNA before further processing. RNA concentration was determined using Qubit RNA HS Assay Kit (#Q32852; Invitrogen). DNA was degraded using the TURBO DNA-free kit (#AM1907; Invitrogen) following the rigorous DNase treatment protocol for the RNA aliquot. cDNA was generated from RNA using the SuperScript III Reverse Transcriptase kit (#18080093; Invitrogen) following the standard protocol. Both positive (*i.e.,* mock bacterial and fungal community (Colovas *et al.* 2022)) and negative (*i.e.,* nuclease-free water) nucleotide extraction controls were included for each batch of nucleic acid extraction, and all controls were processed along with samples.

V4 region 16S rRNA gene amplification and sequencing of DNA and cDNA were performed by the Research Technology Support Facility at Michigan State University (East Lansing, MI, USA) following standard protocols. Amplification via PCR was accomplished using primers targeting the V4 16S rRNA gene region with 515f = GTGCCAGCMGCCGCGGTAA and 806r = GGACTACHVGGGTWTCTAAT (Kozich *et al.* 2013). The PCR mixture consisted of 7.5 µl 2X Dream Taq master mix (#K1071; Thermo Scientific, Waltham USA), 6.5 µl 0.5 µM primer mix, and 1 µl template DNA or cDNA. PCR was performed using the following protocol: 95˚C for 3 min, 30 cycles of 95˚C for 45 s, 50˚C for 60 s, 72˚C for 90 s, and finally 72˚C for 10 min. During amplification, dual indexing barcodes were added, and samples were pooled into two libraries following cleanup. Sequencing was performed on an Illumina MiSeq (Illumina, San Diego USA) with the 2x250 paired-end V2 kit. Triplicate sequencing runs were performed for both DNA libraries, while single runs were performed for both RNA libraries.

To observe overall Bacterial and Archaeal population sizes across the disturbance intensity gradient, 16S rRNA gene counts in undiluted DNA extracts were quantified with qPCR as detailed in supplemental materials. DNA extracts were first diluted 1:20 in PCR grade water. qPCR was performed with triplicate technical replicates using 10 µl SsoAdvanced Universal SYBR Green Supermix (#1725016; BioRad), 1 µl each 10 µM V4 16S rRNA gene primer as before, 6 µl PCR grade water, and 2 µl template DNA solution in 96-well plates. Seven standards were run in triplicate for each 96 well plate based on a 1:4 serial dilution ranging from 72.5 million copies to 17,700 copies per µl using DNA from a culture of *E. coli* MG1655 grown to stationary phase in LB broth and extracted using Quick-DNA Fungal/Bacterial Miniprep kit (#D6005; Zymo). qPCR was performed on the CFX Connect Real-Time PCR Detection System using the following protocol: 95˚C for 15 min, 30 cycles of 94˚C for 45 sec, 50˚C for 60 sec, and 72˚C for 90 sec, followed by a melt curve quality check. Technical replicates were averaged prior to further analysis.

*Sequence processing*

Raw, demultiplexed sequencing reads are available through the NCBI short read archive (BioProject PRJNA973689). Individually for each sequencing run, raw reads were first trimmed using cutadapt (v1.18) (Martin 2011) to remove remaining primer sequences. Paired-end reads were then merged using PEAR (v0.9.6) (Zhang *et al.* 2014) with a maximum length of 600 bp and a minimum overlap of 20 bp. Paired reads with a maximum expected error greater than one and/or containing unknown bases were removed using usearch (v10.0.240_i86linux64) (Edgar 2010). Filtered sequences were aligned to the SILVA SEED reference alignment (v132), and poorly aligned sequences were removed along with those containing homopolymers with a length of 8 or greater using mothur (v1.40.4) (Schloss *et al.* 2009). At this point, quality-filtered sequences from all sequencing runs, including DNA and cDNA libraries, were pooled into a master dataset. Final operational taxonomic units (OTU) were then clustered *de novo* at 97% sequence identity using usearch (Edgar 2010) with chimera removal. Representative OTU sequences in FASTA format are available through figshare (DOI:

10.6084/m9.figshare.23060354). OTU taxonomy was assigned based on the SILVA reference database (v132) using sintax from vsearch (v2.15.2_linux_x86_64) (Rognes *et al.* 2016) with a probability cutoff of 0.8. Following the removal of non-bacterial OTUs (*i.e., Archaea*, mitochondria, chloroplast, and unclassified domain), a phylogenetic tree was generated using FastTree (v2.1.10) (Price *et al.* 2009) with *Sulfolobus* (X90478) included as an outgroup aligned with ssu-align (v0.1.1) (Nawrocki 2009). The sample metadata, OTU table, taxonomy table, and phylogenetic tree were combined into a phyloseq (McMurdie & Holmes 2013) object available through figshare (DOI: 10.6084/m9.figshare.23060354). The sequence processing code is available at https://github.com/ShadeLab/Centralia_RNA_DNA_multiyear.

*Phantom OTUs*

DNA and cDNA sequence set rarefaction produced 50,242 OTUs across both datasets, 38,020 (75.7%) of which were considered active in at least one sample. For each sample, 50.5 ± 0.6% of these active OTUs could be considered phantom (*i.e.,* cDNA reads found, but no reads in the corresponding DNA samples after rarefaction). However, this phantom percentage drops to 35.6 ± 0.7% when considering unrarefied DNA reads and 2.1 ± 0.1% when considering the entire DNA dataset regardless of the sample (Fig. S13), providing evidence that many phantom taxa within the rarefied dataset are likely biological sequences and not entirely artifacts.

*Upset plots*

Upset plots were made using custom code based on the R package UpSetR (Conway *et al.* 2017). For these plots, reference and recovered sites were combined as an overall “no-fire” category because minimal change was observed over time in their active community subsets, and all sites within a year were combined for the new fire-affected and no-fire groups. Thus, an OTU was considered active within a specific year if it was observed as active in at least one site within that year. Only OTUs found to be active in at least five years were included in this analysis.

*Community assembly analysis*

Community assembly processes contributing to the whole bacterial communities' structure were accomplished using NTI calculations (Stegen *et al.* 2012, 2015; Dini-Andreote *et al.* 2015; Barnett *et al.* 2020; Ning *et al.* 2020). Before calculations, the DNA sample OTU table was rarefied to 10,000 reads to reduce computational load. bNTI and $RC_{bray}$ calculations were conducted for site pairs within a year. bNTI was calculated using the big data calculation method from package iCAMP (Ning *et al.* 2020). $RC_{bray}$ was calculated using code modified from Barnett *et al.* 2020 (Barnett *et al.* 2020). bNTI was performed using the full phylogenetic tree, assuming all OTUs recovered are present in some capacity within the regional species pool. In contrast, $RC_{bray}$ used separate metacommunity abundance profiles for each year to account for changes in the structure of the regional species pool.

### 2. Supplemental Figures

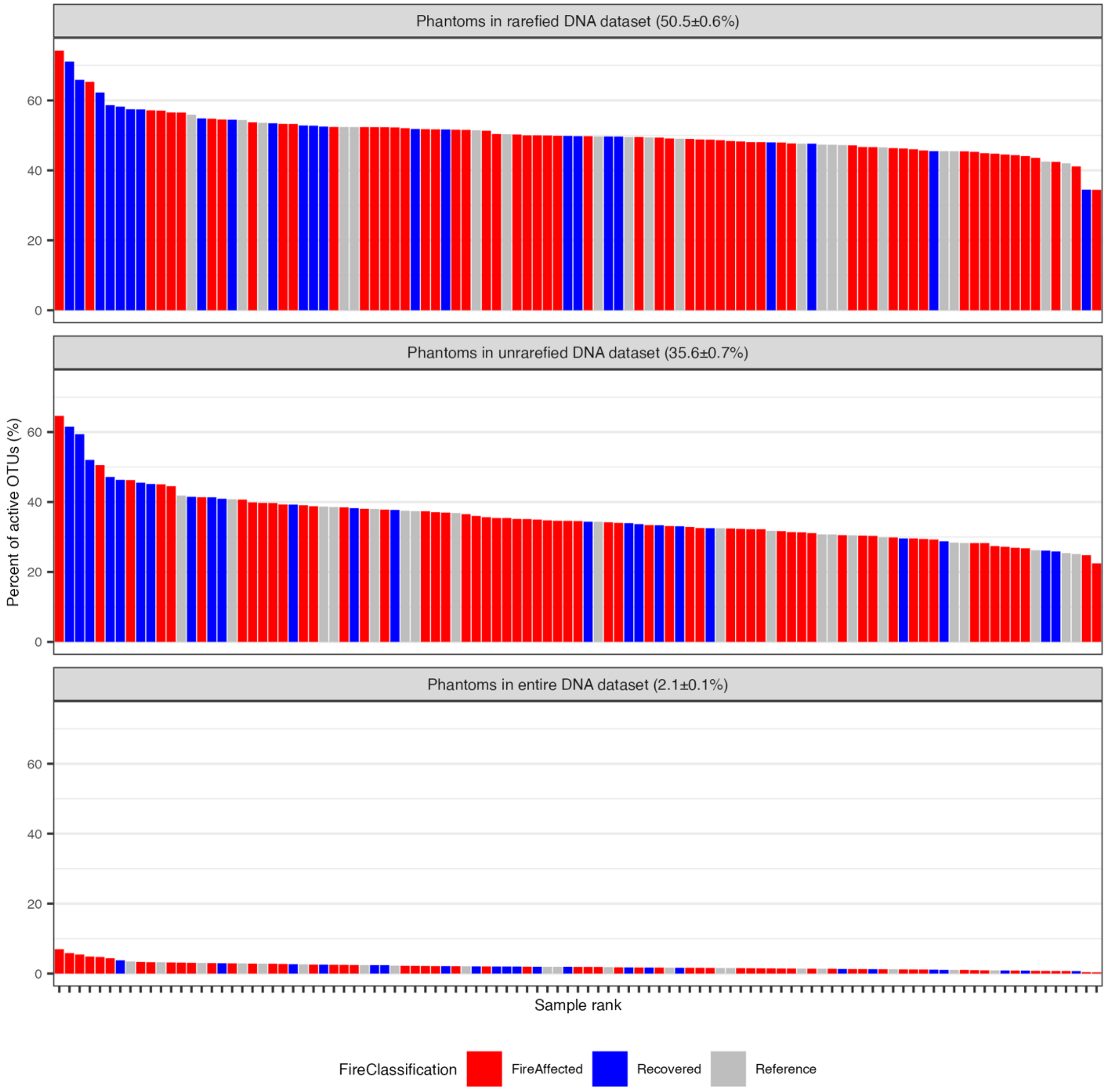


**Figure S1:** Percent of active taxa in each sample that are phantom as defined 3 ways. A) OTUs found in the RNA sample but not in the paired DNA sample where both datasets rarefied to 100,491 reads. B) OTUs found in the RNA sample but not in the paired DNA sample where the DNA dataset is unrarefied. C) OTUs found in the RNA sample but not in any DNA sample regardless of sample pairing and where the DNA dataset is unrarefied.

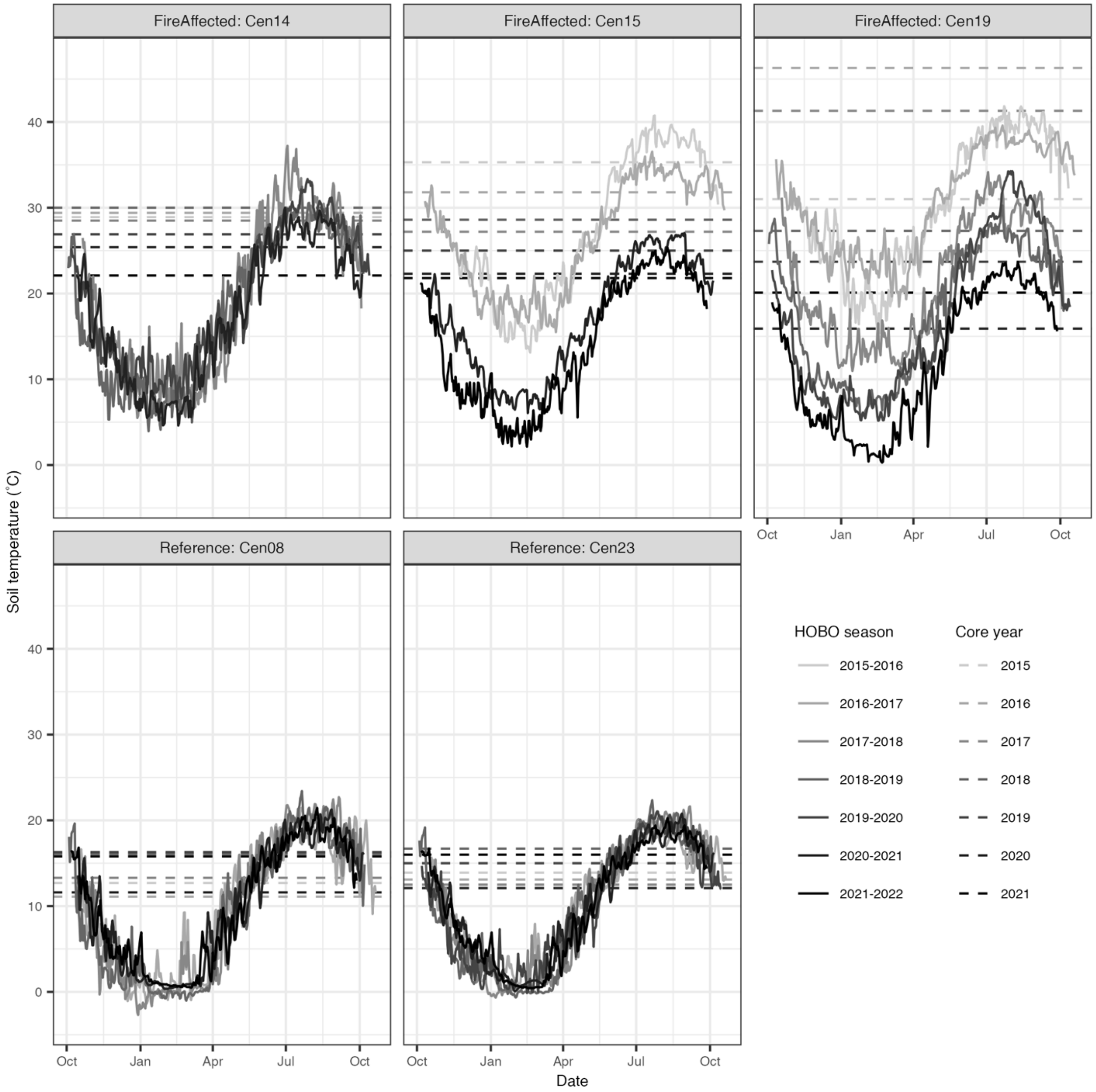


**Figure S2:** Soil temperature data tracked throughout the year for multiple sampling years using the HOBO pendant trackers. Solid lines follow average daily soil temperatures, with color (HOBO season) indicating tracked season (*e.g.,* October 2015-October 2016). Dashed horizontal lines indicate soil core temperatures measured once, just prior to soil core collection with a digital thermometer, with color (Core year) indicating the sampling year.

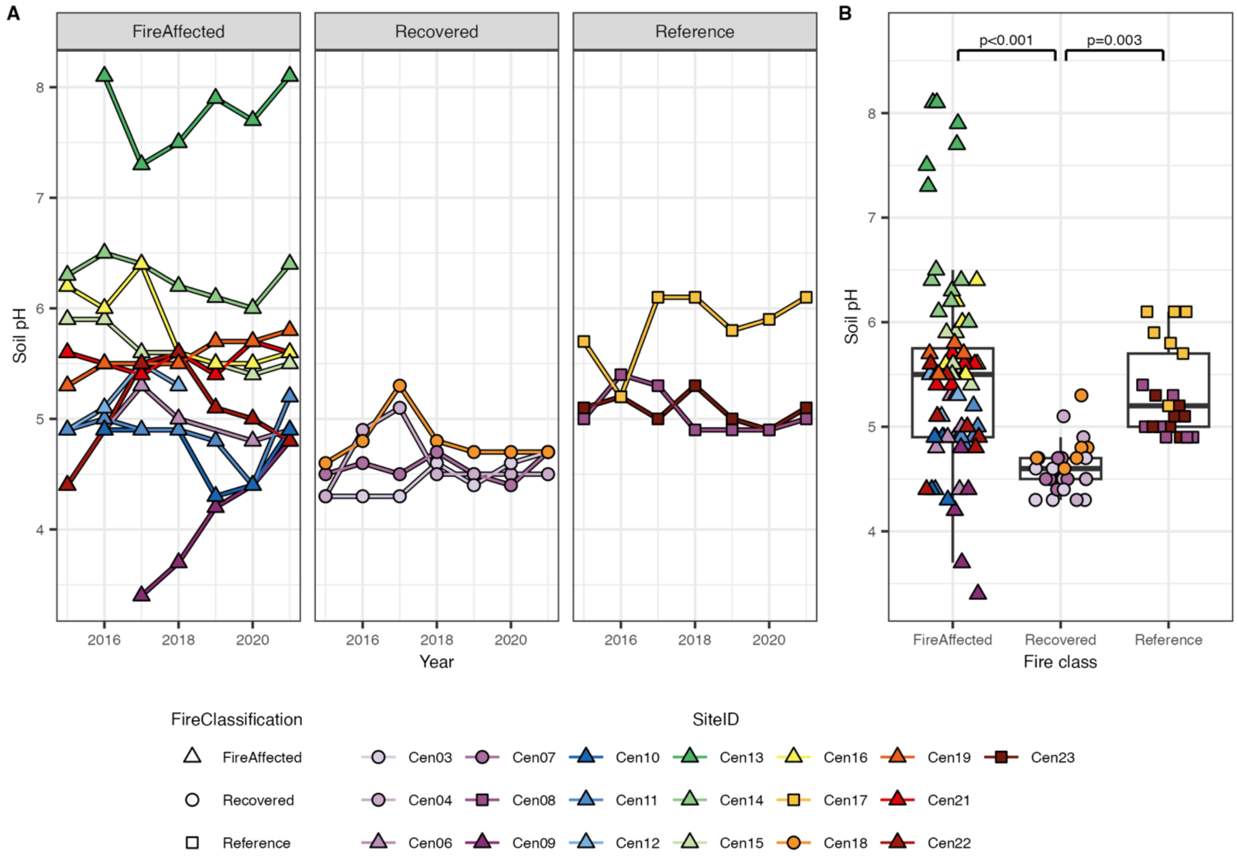


**Figure S3:** Soil pH does not directionally change over time for any fire classification but is generally more acidic in recovered soils than fire affected or reference soils. A) Soil pH in each site over the seven sampling years. No regression slope was statistically significant. B) Difference in overall pH across fire classifications. Post hoc Tukey's test p-values < 0.05 are displayed above statistical significance indicator bars. For all plots, colored points and lines indicate site ID and fire classification.

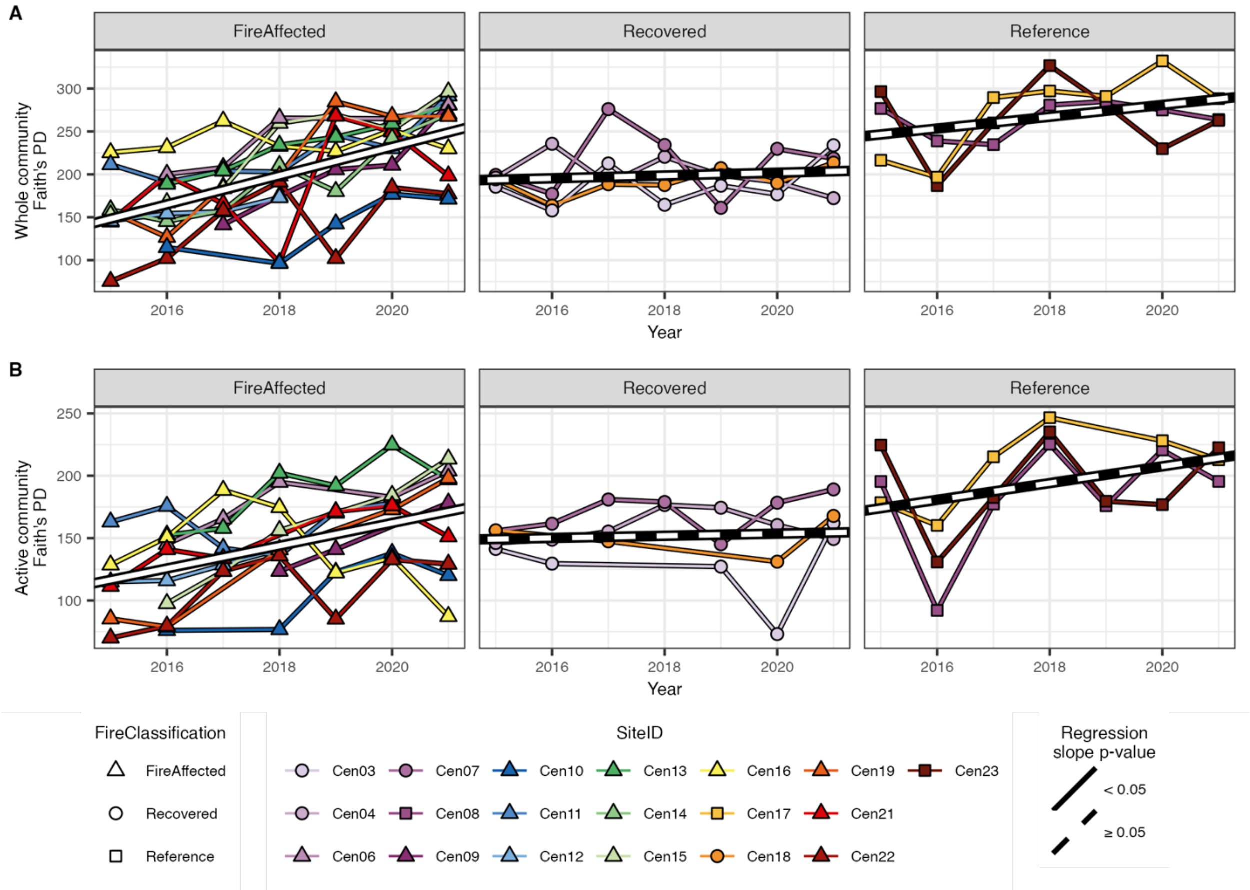


**Figure S4:** Faith's phylogenetic diversity increases over time in the fire affected sites for both whole bacterial communities (A) and active subsets (B). White and black lines indicate linear mixed effects regressions using site ID as the random effect with solid white lines indicating statistically significant slopes (p-value < 0.05) and dashed lines indicating non-statistical significance (p-value ≥ 0.05). For both plots, colored points and lines indicate site ID and fire classification.

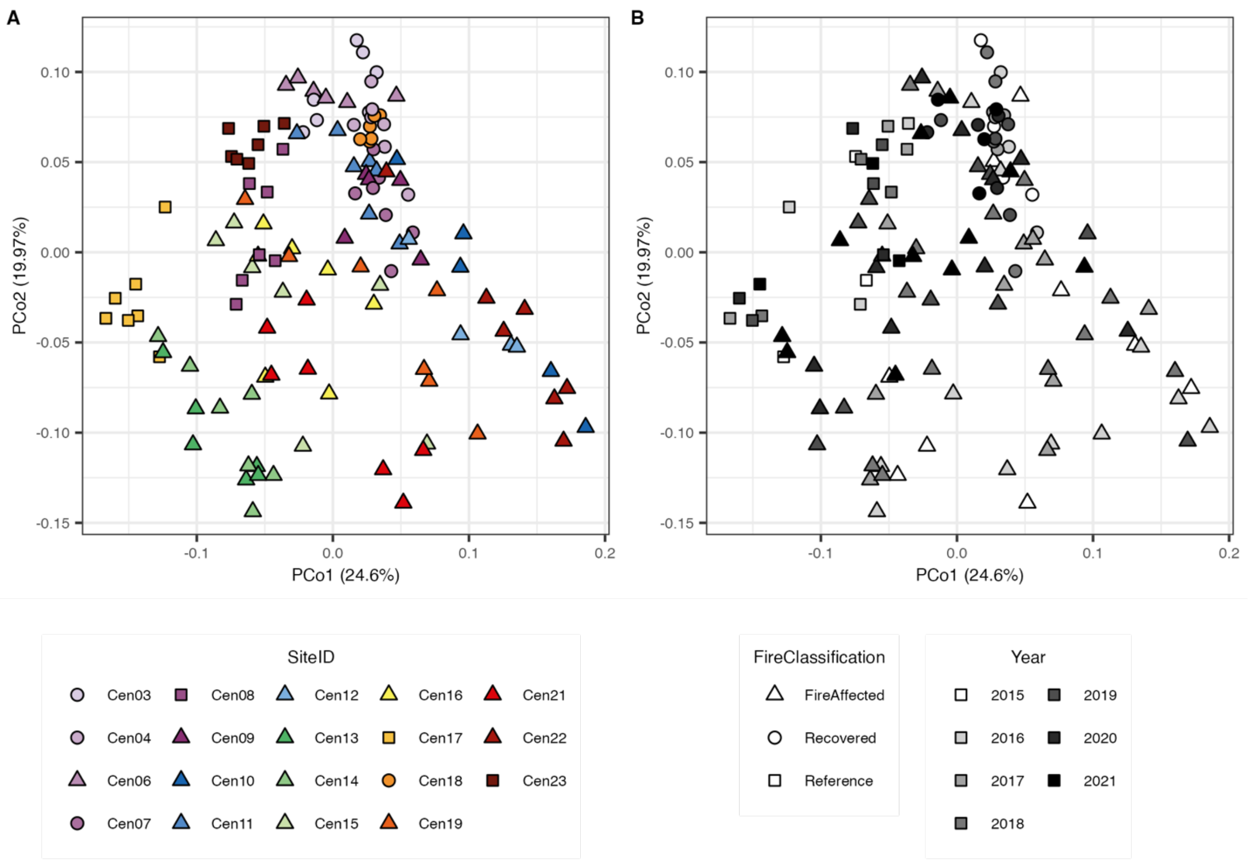


**Figure S5:** Whole bacterial community structure varies across fire classifications and time as demonstrated in principal component analysis (PCoA) based on abundance weighted UniFrac distance. Points in both plots are identical with different color schemes indicating A) site ID and B) sampling year. For both ordinations, shape indicates fire classification.

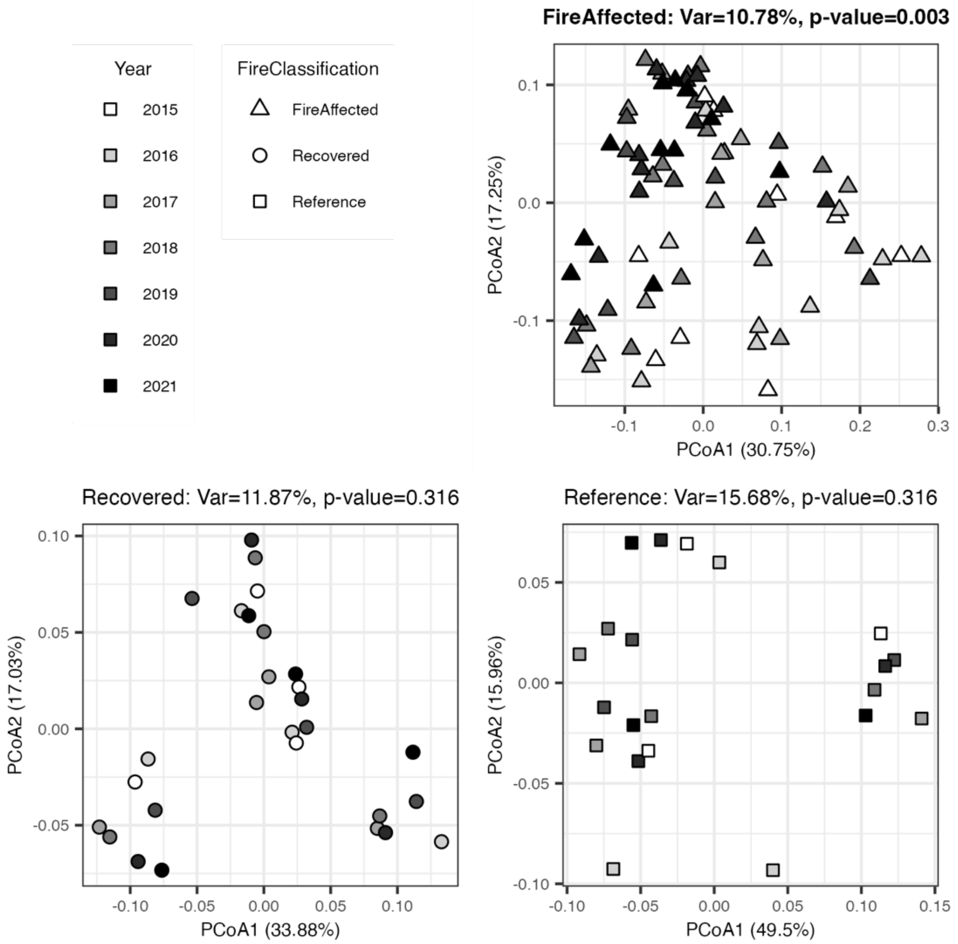


**Figure S6:** Post hoc whole community principal component analysis (PCoA) ordinations examining variation in community structure across sampling years within each fire classification based on abundance weighted UniFrac distance. PERMANOVA results (Table S5) are printed above each point. Time explains a statistically significant portion of variation in community structure only for the fire affected sites.

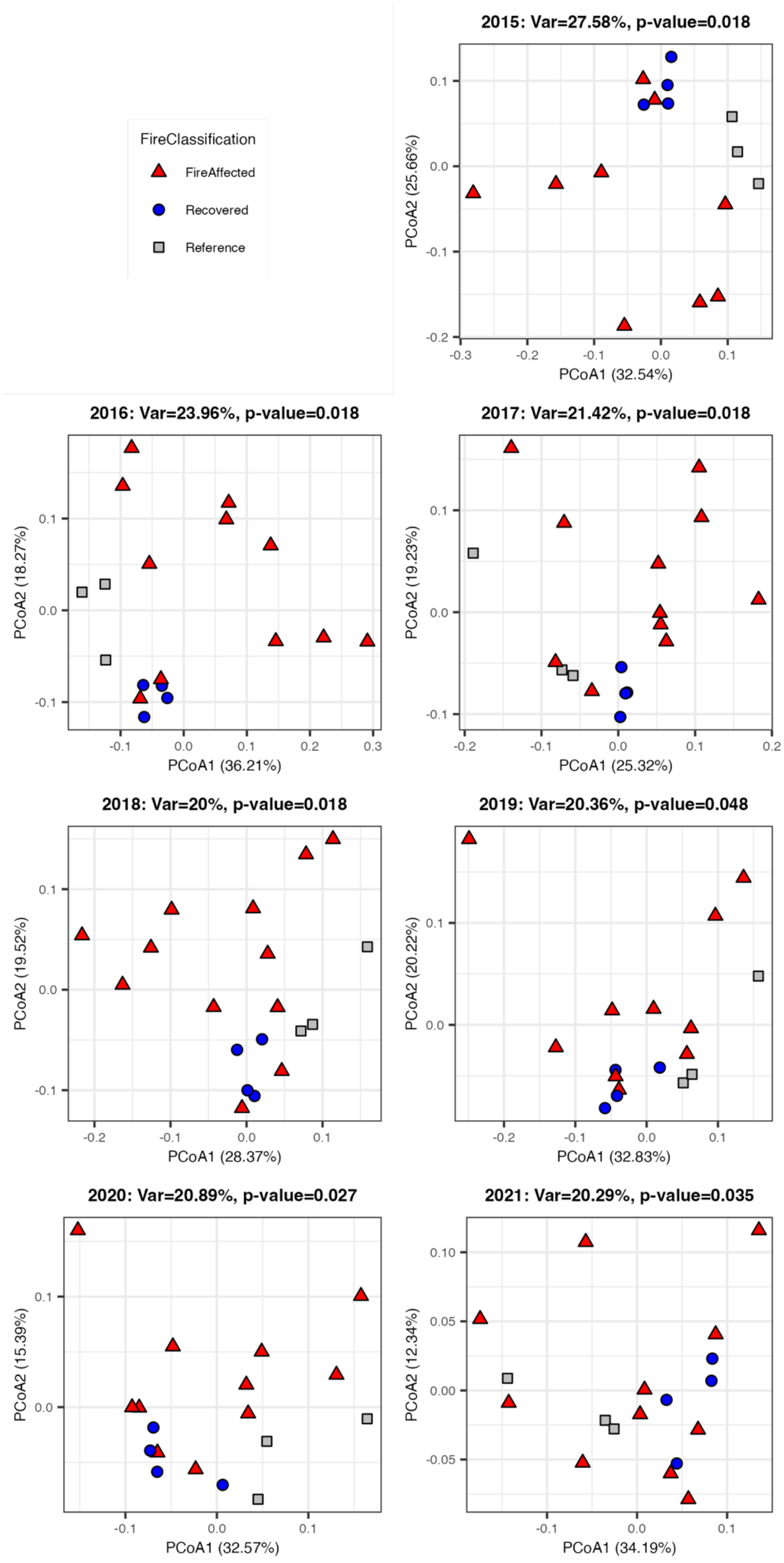
FireClassification
FireAffected
Recovered
Reference
2015: Var=27.58%, p-value=0.018
PCoA1 (32.54%)
PCoA2 (25.66%)
2016: Var=23.96%, p-value=0.018
PCoA1 (36.21%)
PCoA2 (18.27%)
2017: Var=21.42%, p-value=0.018
PCoA1 (25.32%)
PCoA2 (19.23%)
2018: Var=20%, p-value=0.018
PCoA1 (28.37%)
PCoA2 (19.52%)
2019: Var=20.36%, p-value=0.048
PCoA1 (32.83%)
PCoA2 (20.22%)
2020: Var=20.89%, p-value=0.027
PCoA1 (32.57%)
PCoA2 (15.39%)
2021: Var=20.29%, p-value=0.035
PCoA1 (34.19%)
PCoA2 (12.34%)

**Figure S7:** Post hoc whole community principal component analysis (PCoA) ordinations examining variation in community structure across fire classifications within each sampling year based on abundance weighted UniFrac distance. PERMANOVA results (Table S5) are printed above each point. Fire classification explains a statistically significant portion of variation in community structure for all seven years, but the percent of this variation explained decreases over the years.

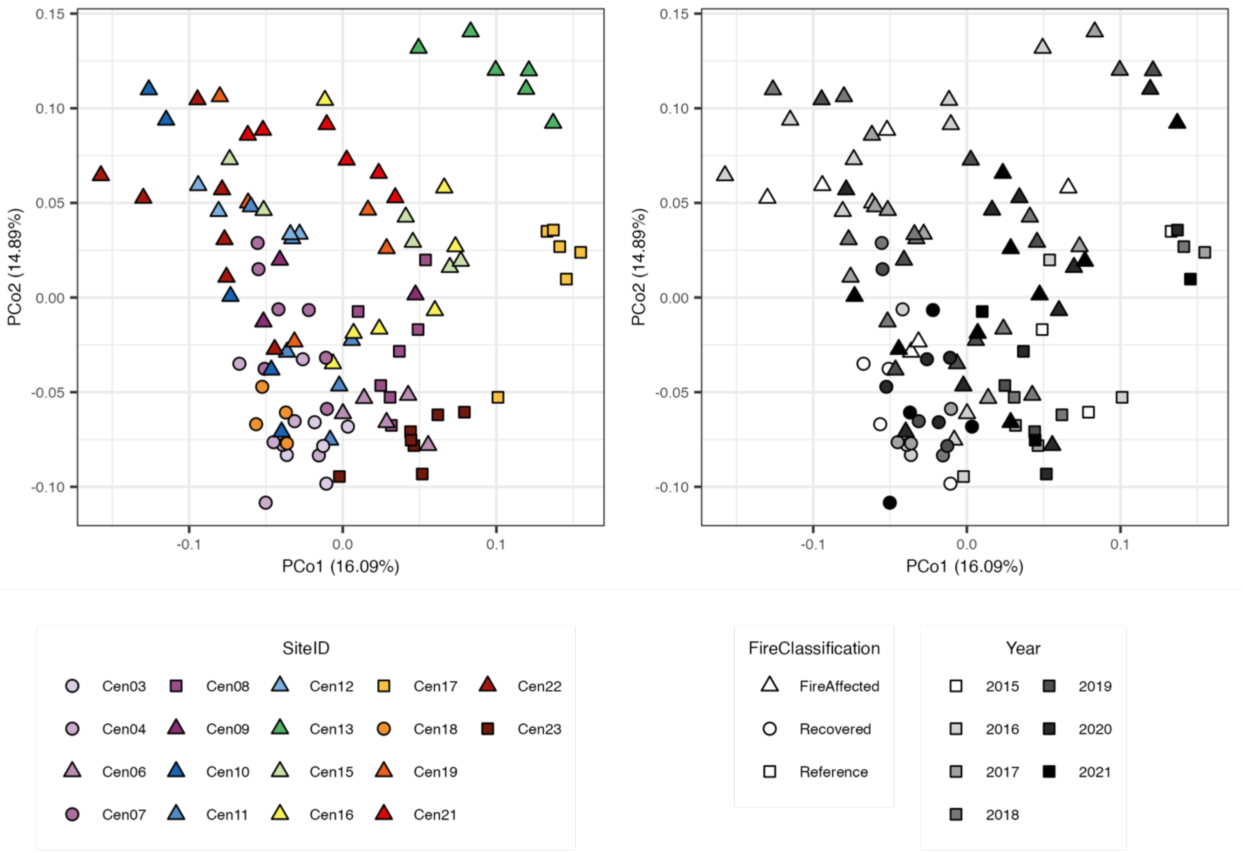


**Figure S8:** Active bacterial community subset structure varies across fire classifications and time as demonstrated in principal component analysis (PCoA) based on abundance weighted UniFrac distance. Points in both plots are identical with different color schemes indicating A) site ID and B) sampling year. For both ordinations, shape indicates fire classification.

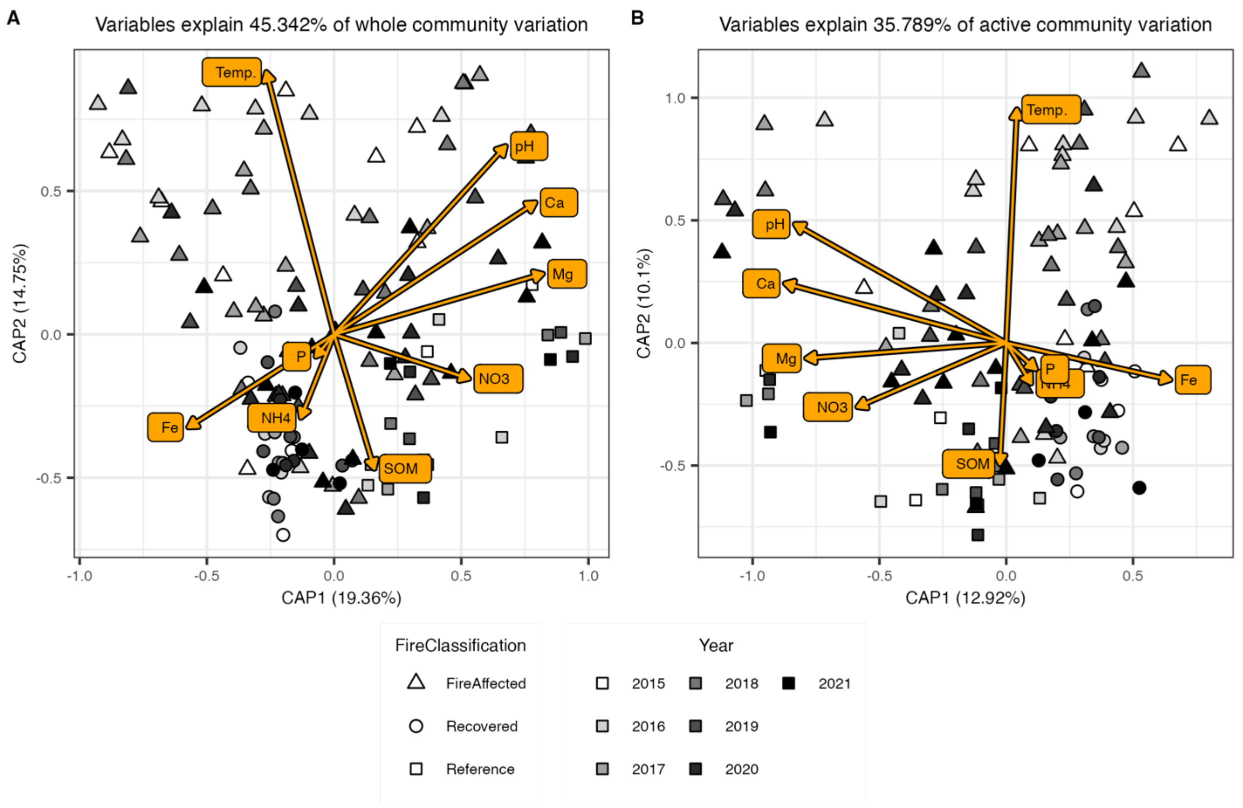


**Figure S9:** Variance in both whole bacterial community (A) and active subset (B) structure is explained by edaphic factors, particularly those associated with temperature and pH. Differences in bacterial community structure is determined with abundance weighted UniFrac and analyzed with constrained correspondence analysis (CCA). Both analyses incorporate soil core temperature (˚C; temp.), pH, calcium (Ca), magnesium (Mg), nitrate ($NO_3$), ammonia ($NH_4$), arsenic (As), potassium (K), phosphate (P), and iron (Fe) concentrations (in ppm), and soil organic matter (SOM) percentage. For both plots, colored points indicate sampling year and fire classification.

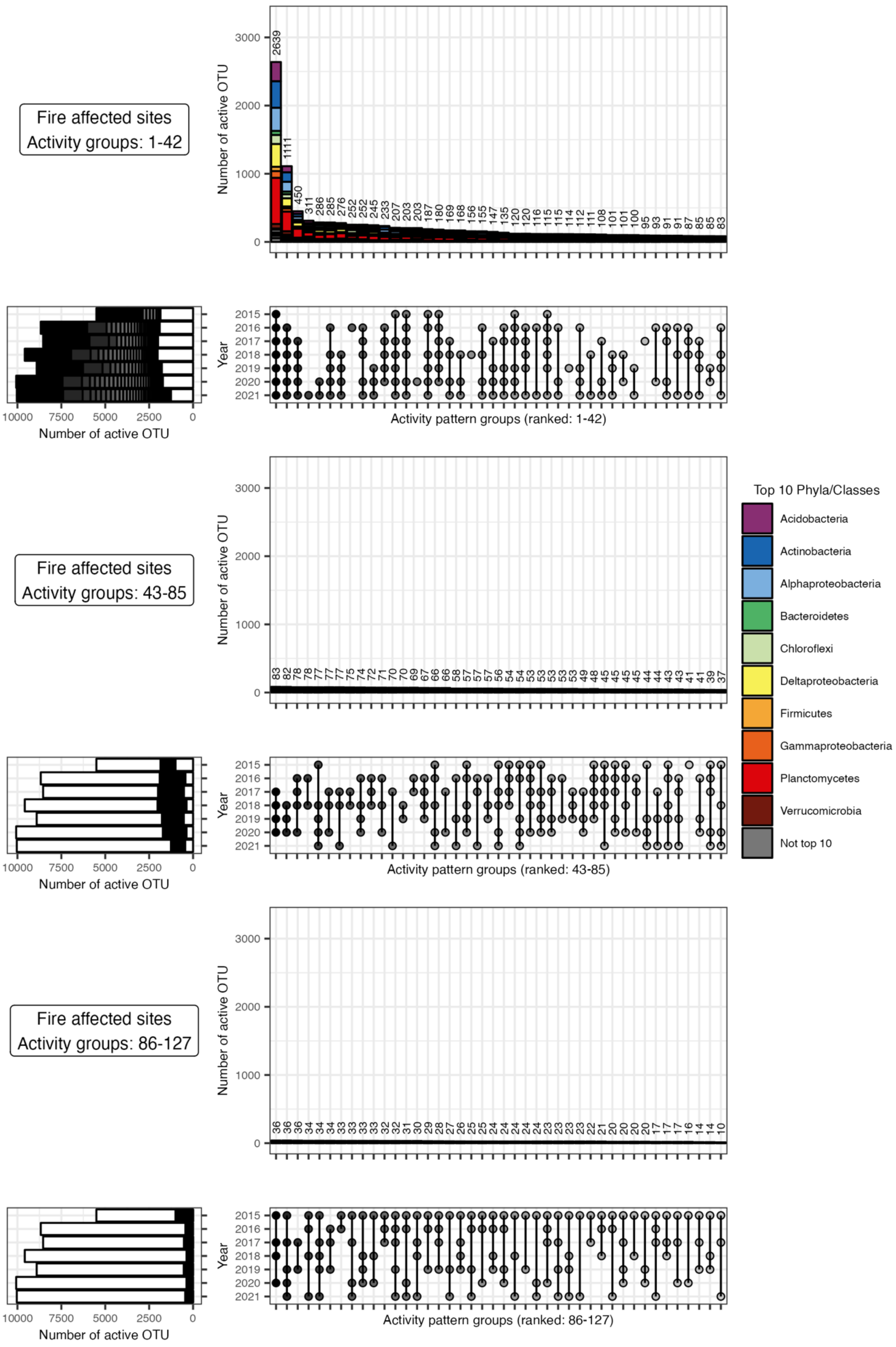

Fire affected sites
Activity groups: 1-42
Number of active OTU
Year
Activity pattern groups (ranked: 1-42)
Fire affected sites
Activity groups: 43-85
Activity pattern groups (ranked: 43-85)
Fire affected sites
Activity groups: 86-127
Activity pattern groups (ranked: 86-127)
Top 10 Phyla/Classes
Acidobacteria
Actinobacteria
Alphaproteobacteria
Bacteroidetes
Chloroflexi
Deltaproteobacteria
Firmicutes
Gammaproteobacteria
Planctomycetes
Verrucomicrobia
Not top 10

**Figure S10:** UpSet plots of all 127 activity across years groups for fire affected sites. All sites sampled within each year were combined such that OTUs were defined as active if they had an RNA:DNA $\geq 1$ in at least one site within that year. Plots are split into three sets of 42-43 groups for better visualization. Groups are ordered by decreasing OTU membership. The bottom right plots show the activity patterns of the listed groups with dots indicating activity within that year and groups ordered by OTU membership. The top vertical bar chart shows the number of the OTUs within each activity group, colored by the top 10 phyla (class for *Proteobacteria*). The bottom left horizontal bar chart shows the number of active OTUs within each year colored by their membership in the listed activity groups. White sections of the bars indicate OTUs from the other unlisted 84-85 groups.

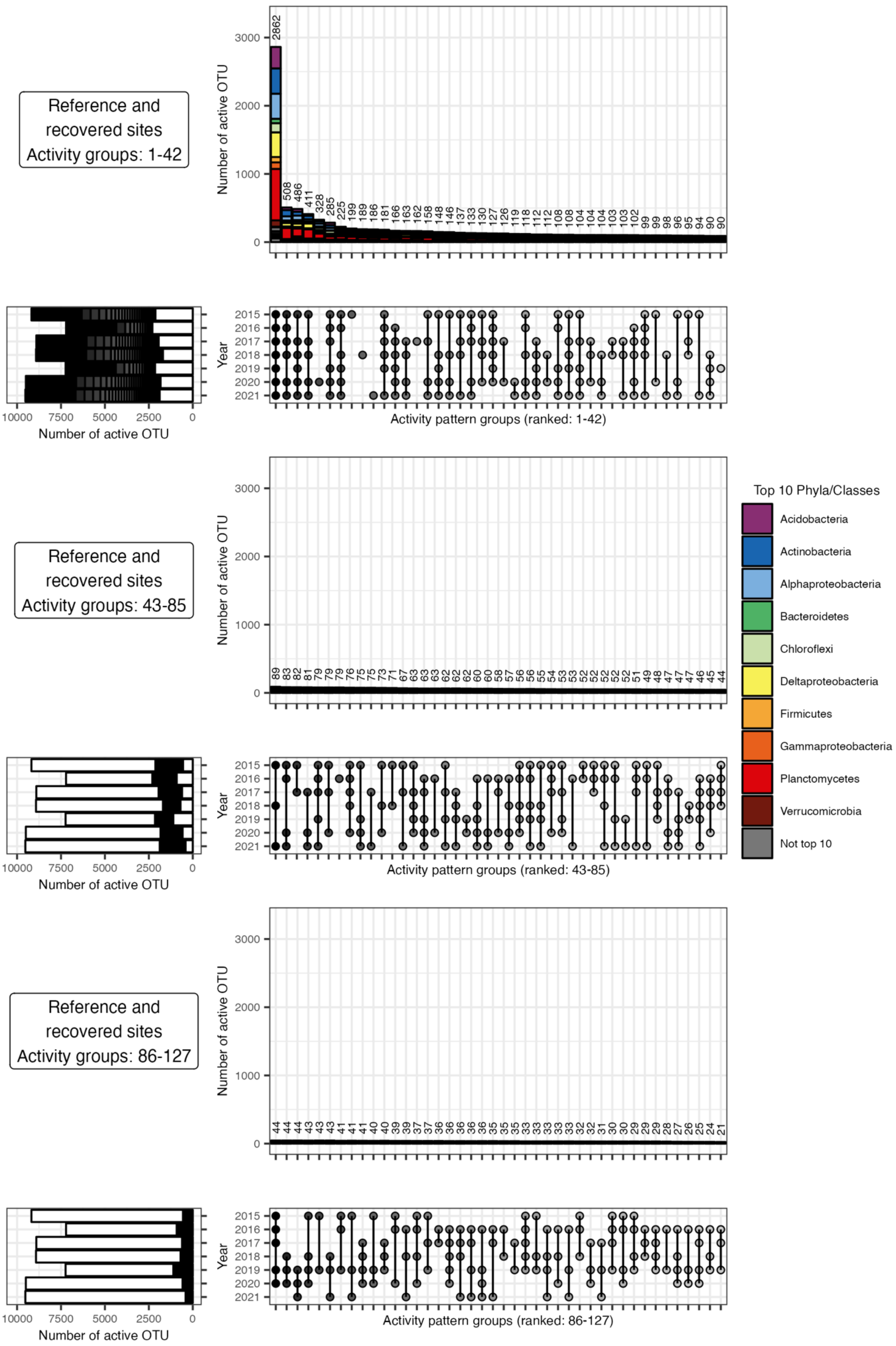
Reference and recovered sites
Activity groups: 1-42
Number of active OTU
Year
2015
2016
2017
2018
2019
2020
2021
10000
7500
5000
2500
0
1000
2000
3000
Activity pattern groups (ranked: 1-42)
Reference and recovered sites
Activity groups: 43-85
Activity pattern groups (ranked: 43-85)
Reference and recovered sites
Activity groups: 86-127
Activity pattern groups (ranked: 86-127)
Top 10 Phyla/Classes
Acidobacteria
Actinobacteria
Alphaproteobacteria
Bacteroidetes
Chloroflexi
Deltaproteobacteria
Firmicutes
Gammaproteobacteria
Planctomycetes
Verrucomicrobia
Not top 10

**Figure S11:** UpSet plots of all 127 activity across years groups for combined reference and recovered sites. All sites sampled within each year were combined such that OTUs were defined as active if they had an RNA:DNA ≥ 1 in at least one site within that year. Plots are split into three sets of 42-43 groups for better visualization. Groups are ordered by decreasing OTU membership. The bottom right plots show the activity patterns of the listed groups with dots indicating activity within that year and groups ordered by OTU membership. The top vertical bar chart shows the number of the OTUs within each activity group, colored by the top 10 phyla (class for *Proteobacteria*). The bottom left horizontal bar chart shows the number of active OTUs within each year colored by their membership in the listed activity groups. White sections of the bars indicate OTUs from the other unlisted 84-85 groups.

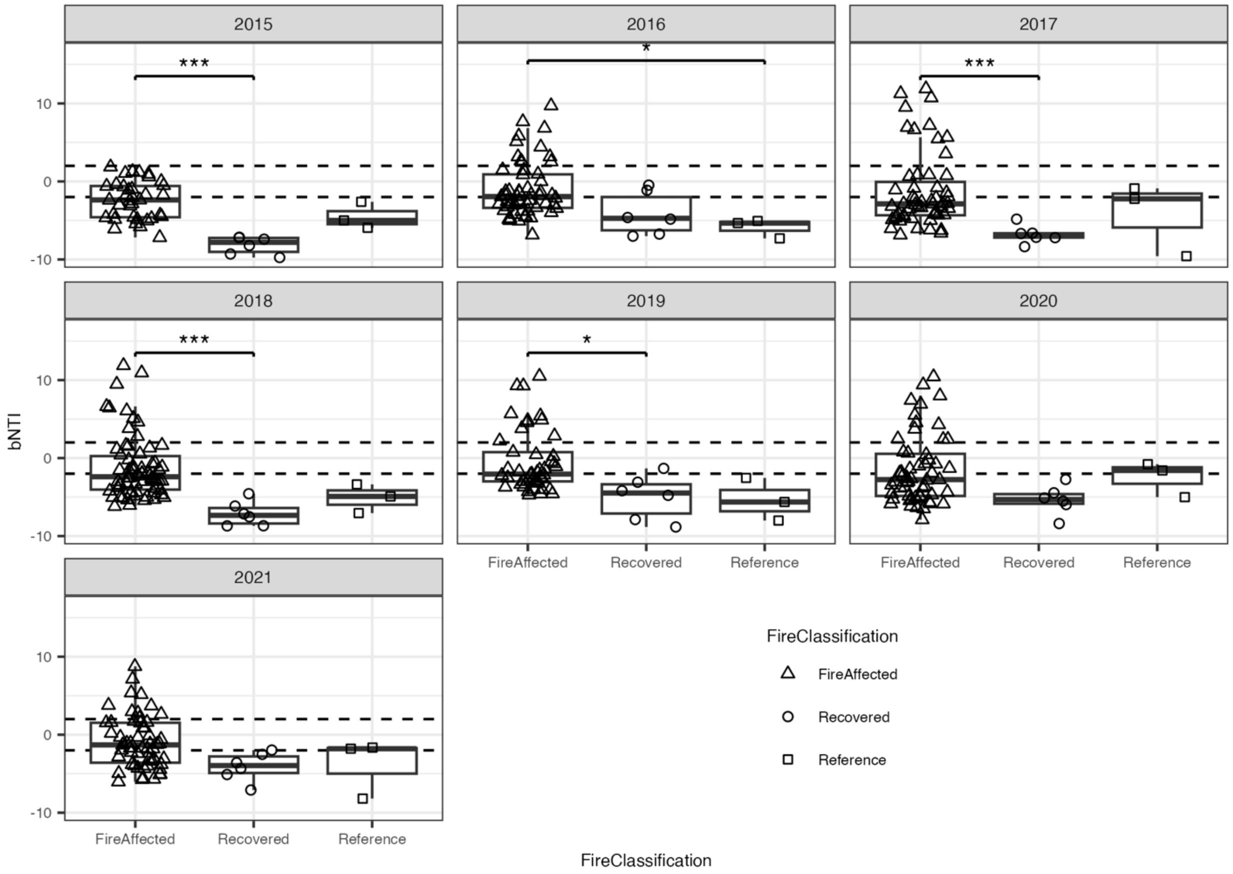


**Figure S12:** bNTI values across fire classifications faceted by sampling year. Similar trends in bNTI are observed across years. Statistically significant *post hoc* pairwise comparisons indicated by significance bars (*: p-value < 0.05, **: p-value < 0.01, ***: p-value < 0.001).

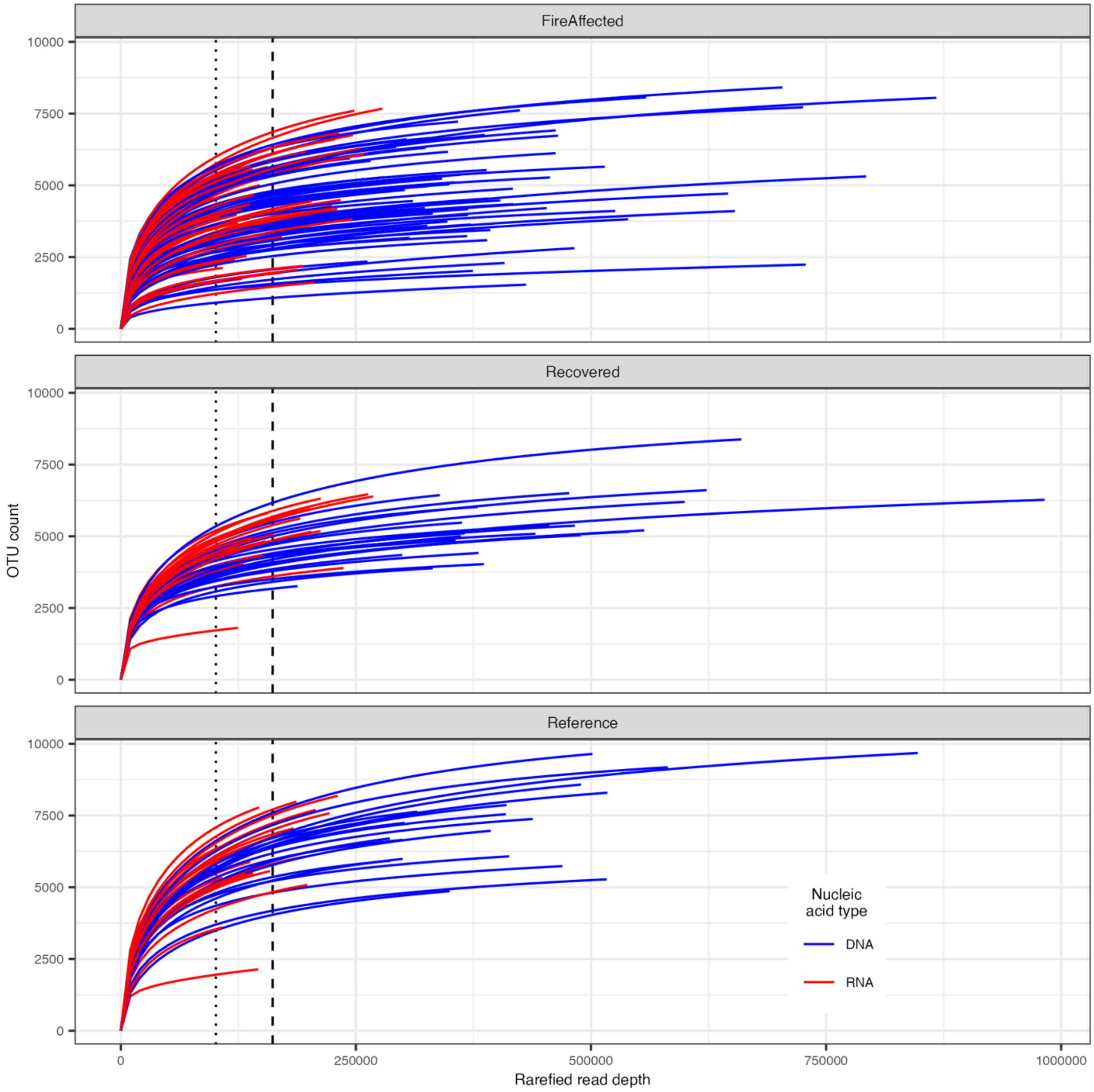


**Figure S13:** Rarefaction curves for both DNA and RNA datasets. Dashed vertical line indicates rarefaction depth for whole community analysis (DNA dataset only; 161,171 reads). Dotted vertical line indicates rarefaction depth for active community analysis (DNA and RNA datasets; 100,491 reads).

### 3. Supplemental Tables

| Edaphic factor | Fire classification | Coefficient | Value | Standard error | Degrees of freedom | t-value | p-value |
|---|---|---|---|---|---|---|---|
| Temperature | FireAffected | Intercept | 3549.521 | 460.582 | 62 | 7.707 | < 0.001 |
| | | Slope | -1.745 | 0.228 | 62 | -7.647 | < 0.001 |
| | Recovered | Intercept | -615.736 | 365.084 | 23 | -1.687 | 0.105 |
| | | Slope | 0.313 | 0.181 | 23 | 1.727 | 0.098 |
| | Reference | Intercept | -639.162 | 409.992 | 17 | -1.559 | 0.137 |
| | | Slope | 0.324 | 0.203 | 17 | 1.594 | 0.129 |
| pH | FireAffected | Intercept | -4.457 | 36.969 | 62 | -0.121 | 0.904 |
| | | Slope | 0.005 | 0.018 | 62 | 0.268 | 0.79 |
| | Recovered | Intercept | -9.807 | 40.492 | 23 | -0.242 | 0.811 |
| | | Slope | 0.007 | 0.02 | 23 | 0.356 | 0.725 |
| | Reference | Intercept | -1.879 | 53.45 | 17 | -0.035 | 0.972 |
| | | Slope | 0.004 | 0.026 | 17 | 0.135 | 0.894 |

**Table S2:** Statistics from linear mixed effects models examining soil temperature and pH change over sampling years. Analysis was done separately for each fire class.

| Community type | Fire Classification | Coefficient | Value | Standard error | Degrees of freedom | t-value | p-value |
|---|---|---|---|---|---|---|---|
| Whole community | FireAffected | Intercept | -33662.431 | 3697.408 | 62 | -9.104 | < 0.001 |
| | | Slope | 16.779 | 1.832 | 62 | 9.158 | < 0.001 |
| | Recovered | Intercept | -3099.864 | 5203.071 | 23 | -0.596 | 0.557 |
| | | Slope | 1.635 | 2.578 | 23 | 0.634 | 0.532 |
| | Reference | Intercept | -13620.816 | 8022.057 | 17 | -1.698 | 0.108 |
| | | Slope | 6.882 | 3.975 | 17 | 1.731 | 0.102 |
| Active community | FireAffected | Intercept | -18270.151 | 3497.704 | 48 | -5.223 | < 0.001 |
| | | Slope | 9.125 | 1.733 | 48 | 5.265 | < 0.001 |
| | Recovered | Intercept | -1589.509 | 3983.957 | 18 | -0.399 | 0.695 |
| | | Slope | 0.863 | 1.974 | 18 | 0.437 | 0.667 |
| | Reference | Intercept | -13345.299 | 7943.471 | 16 | -1.68 | 0.112 |
| | | Slope | 6.709 | 3.936 | 16 | 1.704 | 0.108 |

**Table S3:** Statistics from linear mixed effects models examining bacterial community phylogenetic diversity change over sampling years. Analysis was done separately for each fire class.

| Community type | Coefficient | Value | Standard error | Degrees of freedom | t-value | p-value |
|---|---|---|---|---|---|---|
| Whole community | Intercept | 297.193 | 16.633 | 104 | 17.868 | < 0.001 |
| | Slope | -3.853 | 0.652 | 104 | -5.908 | < 0.001 |
| Active community | Intercept | 204.492 | 13.364 | 84 | 15.302 | < 0.001 |
| | Slope | -2.227 | 0.538 | 84 | -4.137 | < 0.001 |

**Table S4:** Statistics from linear mixed effects models examining bacterial community phylogenetic diversity relationship to soil core temperature. Analysis was done with all fire classifications combined.

| Coefficient | Value | Standard error | Degrees of freedom | t-value | p-value |
|---|---|---|---|---|---|
| Intercept | 5150000000 | 1030000000 | 101 | 4.985 | 2.58E-06 |
| Slope | -76000000 | 40500000 | 101 | -1.877 | 6.34E-02 |

**Table S5:** Statistics from linear mixed effects models examining microbial count relationship to soil core temperature. Analysis was done with all fire classifications combined.

| Community type | Factor | Degrees of freedom | Sum of Squares | R2 | F value | p-value |
|---|---|---|---|---|---|---|
| Whole community | Fire Classification | 2 | 0.497 | 0.18 | 12.811 | 0.001 |
| | Year | 6 | 0.157 | 0.057 | 1.35 | 0.001 |
| | Fire Classification: Year | 12 | 0.11 | 0.04 | 0.474 | 0.002 |
| | Residual | 103 | 1.997 | 0.723 | | |
| | Total | 123 | 2.762 | 1 | | |
| Active community | Fire Classification | 2 | 0.343 | 0.124 | 6.793 | 0.001 |
| | Year | 6 | 0.158 | 0.057 | 1.042 | 0.002 |
| | Fire Classification: Year | 12 | 0.184 | 0.067 | 0.608 | 0.18 |
| | Residual | 82 | 2.068 | 0.751 | | |
| | Total | 102 | 2.753 | 1 | | |

**Table S6:** Statistics from PERMANOVA analyses on abundance weighted UniFrac distance blocked by site ID for whole bacterial communities and active subsets.

| Fire classification | Year | Factor | Degrees of freedom | Sum of Squares | R2 | F value | p-value | Adjusted p-value |
|---|---|---|---|---|---|---|---|---|
| NA | 2015 | FireClassification | 2 | 0.155 | 0.276 | 2.476 | 0.005 | 0.018 |
| | 2016 | | 2 | 0.182 | 0.24 | 2.364 | 0.01 | 0.018 |
| | 2017 | | 2 | 0.123 | 0.214 | 2.045 | 0.006 | 0.018 |
| | 2018 | | 2 | 0.115 | 0.2 | 2.001 | 0.01 | 0.018 |
| | 2019 | | 2 | 0.098 | 0.204 | 1.79 | 0.048 | 0.048 |
| | 2020 | | 2 | 0.089 | 0.209 | 1.98 | 0.019 | 0.027 |
| | 2021 | | 2 | 0.079 | 0.203 | 1.91 | 0.03 | 0.035 |
| FireAffected | NA | Year | 6 | 0.294 | 0.108 | 1.369 | 0.001 | 0.003 |
| Recovered | | | 6 | 0.049 | 0.119 | 0.472 | 0.28 | 0.316 |
| Reference | | | 6 | 0.042 | 0.157 | 0.434 | 0.316 | 0.316 |

**Table S7:** Statistics from post hoc PERMANOVA analyses on abundance weighted UniFrac distance blocked by site ID for whole bacterial communities separately either across years (measuring variability explained by fire classification) or across fire classifications (measuring varaibility explained by year). p-vaues are adjusted using the Benjamini-Hochberg procedure (n=7 and n=3 respectively).

| Community type | Comparison | Difference | Lower endpoint | Upper endpoint | Adjusted p-value |
|---|---|---|---|---|---|
| Whole community | Recovered-FireAffected | -0.054 | -0.073 | -0.035 | < 0.001 |
| | Reference-FireAffected | -0.058 | -0.08 | -0.037 | < 0.001 |
| | Reference-Recovered | -0.004 | -0.029 | 0.021 | 0.921 |
| Active community | Recovered-FireAffected | -0.04 | -0.062 | -0.018 | < 0.001 |
| | Reference-FireAffected | -0.049 | -0.072 | -0.026 | < 0.001 |
| | Reference-Recovered | -0.009 | -0.037 | 0.018 | 0.716 |

**Table S8:** Statistics from post hoc Tukey-HSD test on homogenetic of multivariate dispersons based on abundance weighted UniFrac distance across sites.

| Community type | Fire Classification | Coefficient | Value | Standard error | Degrees of freedom | t-value | p-value |
|---|---|---|---|---|---|---|---|
| Whole community | FireAffected | Intercept | 0.062 | 0.012 | 190 | 5.184 | < 0.001 |
| | | Slope | 0.058 | 0.006 | 190 | 9.661 | < 0.001 |
| | Recovered | Intercept | 0.087 | 0.011 | 79 | 7.93 | < 0.001 |
| | | Slope | 0.005 | 0.004 | 79 | 1.295 | 0.199 |
| | Reference | Intercept | 0.084 | 0.015 | 59 | 5.561 | < 0.001 |
| | | Slope | 0.005 | 0.009 | 59 | 0.606 | 0.547 |
| Active community | FireAffected | Intercept | 0.12 | 0.017 | 129 | 7.004 | < 0.001 |
| | | Slope | 0.047 | 0.009 | 129 | 5.305 | < 0.001 |
| | Recovered | Intercept | 0.143 | 0.022 | 53 | 6.453 | < 0.001 |
| | | Slope | 0.001 | 0.009 | 53 | 0.106 | 0.916 |
| | Reference | Intercept | 0.145 | 0.023 | 53 | 6.202 | < 0.001 |
| | | Slope | -0.004 | 0.014 | 53 | -0.284 | 0.778 |

**Table S9:** Time-lag analysis statistics from linear mixed effects models examining abundance weighted UniFrac distance between years of the same site and the square root of the difference in time. Analysis was done separately for each fire classification.

| Fire Classification | Coefficient | Value | Standard error | Degrees of freedom | t-value | p-value |
|---|---|---|---|---|---|---|
| FireAffected | Intercept | 44.832 | 1.618 | 129 | 27.707 | < 0.001 |
| | Slope | -1.868 | 0.324 | 129 | -5.765 | < 0.001 |
| Recovered | Intercept | 43.847 | 1.947 | 53 | 22.517 | < 0.001 |
| | Slope | -0.516 | 0.349 | 53 | -1.478 | 0.145 |
| Reference | Intercept | 44.857 | 1.423 | 53 | 31.524 | < 0.001 |
| | Slope | -0.134 | 0.383 | 53 | -0.35 | 0.727 |

**Table S10:** Time-lag analysis of percentage of Bray-Curtis dissimilarity attributable to active taxa. Linear mixed effects models examining percentage of Bray-Curtis dissimiliarty between years of the same site and the difference in time. Analysis was done separately for each fire classification.

| Coefficient | Estimate | Standard error | t-value | p-value |
|---|---|---|---|---|
| Intercept | -1.917 | 0.085 | -22.681 | < 0.001 |
| Scaled difference in soil pH | 2.629 | 0.091 | 29.028 | < 0.001 |
| Scaled difference in soil temperature | 0.416 | 0.085 | 4.864 | < 0.001 |
| Scaled difference in soil pH: Scaled difference in soil temperature | -0.096 | 0.072 | -1.335 | 0.182 |

**Table S11:** Statistics for linear regression of scaled and centered differences in soil pH and soil temperature to bNTI for whole bacterial communities.
